# Hepatitis C Virus Protease Inhibitors Show Differential Efficacy and Interactions with Remdesivir for Treatment of SARS-CoV-2 *in Vitro*

**DOI:** 10.1101/2020.12.02.408112

**Authors:** Karen A. Gammeltoft, Yuyong Zhou, Andrea Galli, Anna Offersgaard, Long V. Pham, Ulrik Fahnøe, Shan Feng, Santseharay Ramirez, Jens Bukh, Judith M. Gottwein

## Abstract

Antivirals targeting SARS-CoV-2 could improve treatment of COVID-19. We evaluated the efficacy of clinically relevant hepatitis C virus (HCV) NS3 protease inhibitors (PI) against SARS-CoV-2 and their interactions with remdesivir, the only antiviral approved for treatment of COVID-19. HCV PI showed differential potency in VeroE6 cell-based antiviral assays based on detection of the SARS-CoV-2 Spike protein. Linear PI boceprevir, telaprevir and narlaprevir had 50% effective concentrations (EC50) of ~40 μM. Among macrocyclic PI simeprevir, paritaprevir, grazoprevir, glecaprevir, voxilaprevir, vaniprevir, danoprevir and deldeprevir, simeprevir had the highest (EC50 15 μM) and glecaprevir the lowest (EC50 >178 μM) potency. Acyclic PI asunaprevir and faldaprevir had EC50 of 72 and 23 μM, respectively. ACH-806, an HCV NS3 protease co-factor NS4A inhibitor, had EC50 of 46 μM. For selected PI, potency was similar in human hepatoma Huh7.5 cells. Selectivity indexes, based on antiviral and cell viability assays, were highest for linear PI. In combination with remdesivir, linear PI boceprevir and narlaprevir showed antagonism, while macrocyclic PI simeprevir, paritaprevir and grazoprevir showed synergism with drug reduction indexes of up to 27 for simeprevir. Treatment of infected cultures with equipotent concentrations (1-fold EC50) of HCV PI revealed minor differences in barrier to SARS-CoV-2 escape. Complete viral suppression was achieved treating with ≥3-fold EC50 boceprevir or combination of 1-fold EC50 simeprevir with 0.4-fold EC50 remdesivir, not leading to significant viral suppression in single treatments. Considering potency, human plasma concentrations and synergism with remdesivir, simeprevir seemed the most promising compound for optimization of future antiviral treatments of COVID-19.

## Introduction

Severe acute respiratory syndrome coronavirus 2 (SARS-CoV-2) is a positive-sense single-stranded RNA virus of the *Coronaviridae* family, which emerged in humans in 2019 most likely originating from a bat-borne virus.^1–3^ SARS-CoV-2 causes coronavirus disease 2019 (COVID-19), a multi-systemic disease with initial symptoms mostly localizing to the respiratory tract. Until the end of November 2020, the COVID-19 pandemic has been responsible for >60 million infected, >1.4 million deaths, and an unknown number of individuals suffering from long-term health effects.^4–8^ Repurposing of drugs approved for other medical indications is promoted as a time-saving approach to identification of urgently needed treatments. At present, the only drug approved for treatment of COVID-19 that directly targets SARS-CoV-2 proteins is remdesivir, an inhibitor of the viral nonstructural protein (nsp) 12 polymerase, originally being an investigational broad-spectrum antiviral previously evaluated for treatment of chronic hepatitis C virus (HCV) infection and ebola infection.^9^

Another important target of antiviral drugs are viral proteases, which are essential for cleavage of viral polyproteins into functional proteins.^10–12^ The coronavirus main protease (M^pro^) or 3 chymotrypsin-like protease (3CLpro) is a cysteine protease corresponding to nsp5. M^pro^ mediates 11 cleavage events at conserved sites of the polyprotein and is thus essential for viral replication.^13–15^ In addition, M^pro^ has no cellular homologues and is highly conserved between different coronaviruses, making it an interesting drug target.^16^

Hepatitis C virus is a positive-sense single-stranded RNA virus of the *Flaviviridae* family, which was classified into 8 major genotypes and various subtypes.^17,18^ The main HCV protease, nonstructural protein 3 (NS3), is a chymotrypsin-like serine protease.^19–21^ Together with its essential co-factor, NS4A, it mediates 4 cleavage events of the polyprotein. Inhibitors of this protease are important components of recently developed highly efficient HCV treatment regimens based on combination of antivirals directly targeting HCV proteins.^22^

Initially developed HCV protease inhibitors (PI) showed a linear structure and included boceprevir and telaprevir, which were approved in 2011 in the U.S. and EU, as well as narlaprevir, approved in 2016 in Russia, for treatment of chronic HCV infection (Supplementary Figure 1, Supplementary Table 1). Subsequently, PI with macrocyclic structure, including simeprevir, paritaprevir, grazoprevir, glecaprevir, voxilaprevir, vaniprevir, danoprevir and deldeprevir were developed. These macrocyclic PI were approved between 2013 and 2019 in the U.S., EU or China, with exception of vaniprevir, only approved in Japan and deldeprevir, which was never approved. Of the 2 acyclic PI, asunaprevir and faldaprevir, asunaprevir was approved in Japan, Canada and China, while faldaprevir was not approved. Several of the initially developed PI were subsequently discontinued due to the development of more efficient PI with increased activity against the different HCV genotypes (Supplementary Table 1). At present clinical use in the U.S., Europe and China is focused on inhibitor combinations, including grazoprevir, glecaprevir and voxilaprevir.

Additionally, in China, inhibitor combinations including paritaprevir, danoprevir or asunaprevir are used in the clinic.

While an inhibitor of HCV NS4A (ACH-806) was tested in clinical phase 1 trials, development was halted due to reversible nephrotoxicity.^23,24^

In this study we investigated *in vitro* efficacy of a panel of HCV PI, including all clinically approved compounds and selected compounds tested in clinical studies, against SARS-CoV-2. We further evaluated efficacy of an HCV NS4A inhibitor. In concentration-response antiviral assays we determined 50% effective concentrations (EC50), 50% cytotoxic concentrations (CC50) and selectivity indexes (SI). Moreover, we evaluated interactions with remdesivir for selected linear and macrocyclic compounds. Finally, in longer-term cultures we evaluated selected inhibitors for their barrier to viral escape.

## Results

### Differential potency of clinically relevant HCV protease inhibitors and potency of an HCV NS4A inhibitor against SARS-CoV-2 *in vitro*

To determine the potency of a panel of HCV PI and an HCV NS4A inhibitor against SARS-CoV-2, we developed a cell-based antiviral assay in 96-well plates, adapting an assay previously developed to determine potency of HCV PI against HCV.^24–31^ First, concentration response studies were carried out in VeroE6 cells using inhibitor concentrations not resulting in reduction of cell viability (relative cell viability >90%). All tested inhibitors were able to inhibit the virus with EC50 values in the micromolar range, with exception of glecaprevir, voxilaprevir and deldeprevir, where EC50 values could not be determined due to cytotoxicity of the drugs or antiviral activity of the diluent dimethyl sulfoxide (DMSO) at high drug concentrations (Figure 1, Table 1, Supplementary Figure 2). The linear PI boceprevir, telaprevir and narlaprevir showed comparable potencies with EC50 values of ̴ 40 μM. Among the macrocyclic PI, simeprevir showed the highest potency with an EC50 of 15 μM. Further, paritaprevir had an EC50 of 22 μM, while grazoprevir and vaniprevir had EC50 values of 42 and 51 μM, respectively. Finally, EC50 was 87 μM for danoprevir. For the acyclic PI, faldaprevir (EC50 23 μM) was more potent than asunaprevir (EC50 72 μM).

**Table 1.**
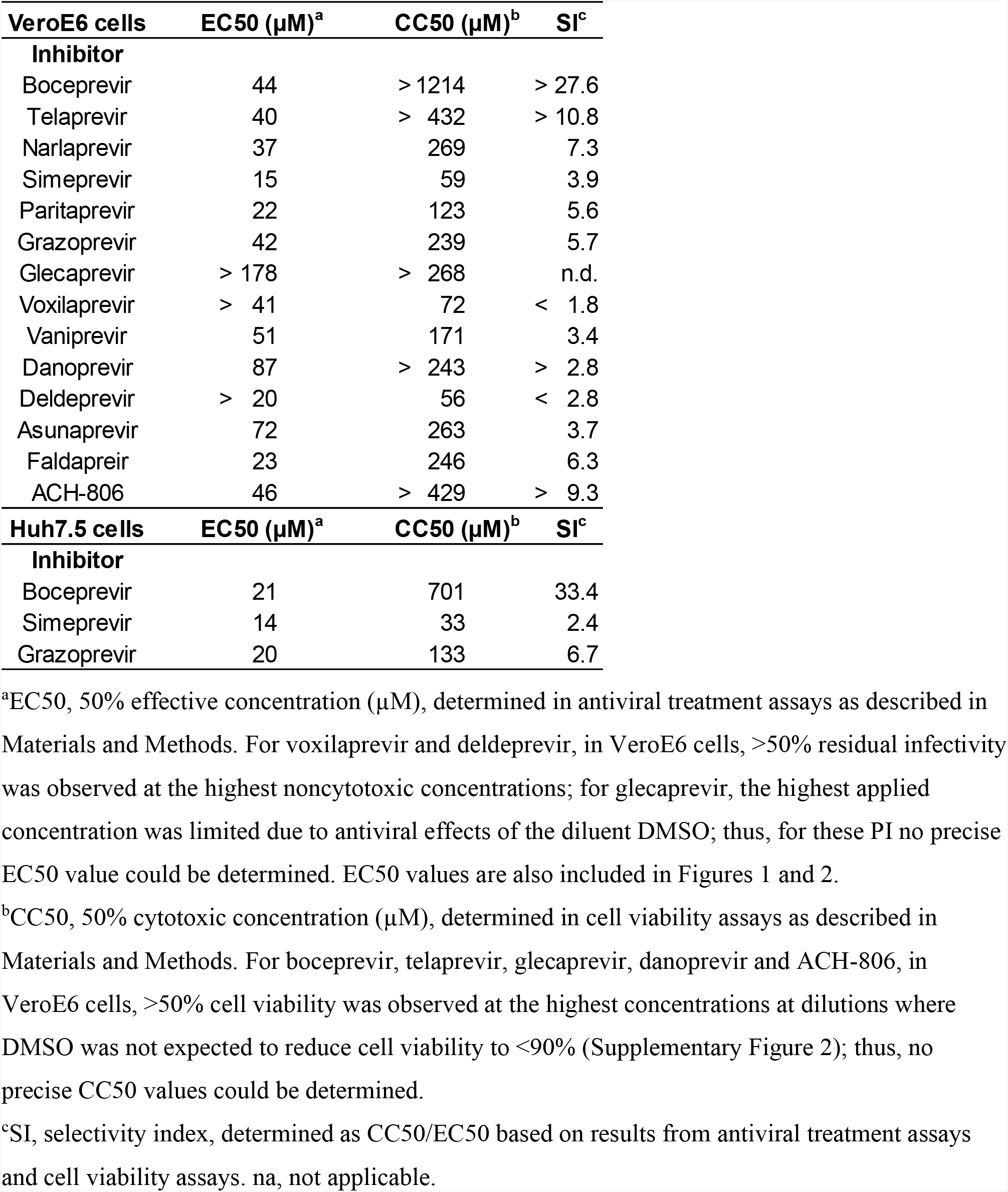
Potency of a panel of HCV PI and an HCV NS4A inhibitor against SARS-CoV-2 *in vitro*.

**Figure 1.**
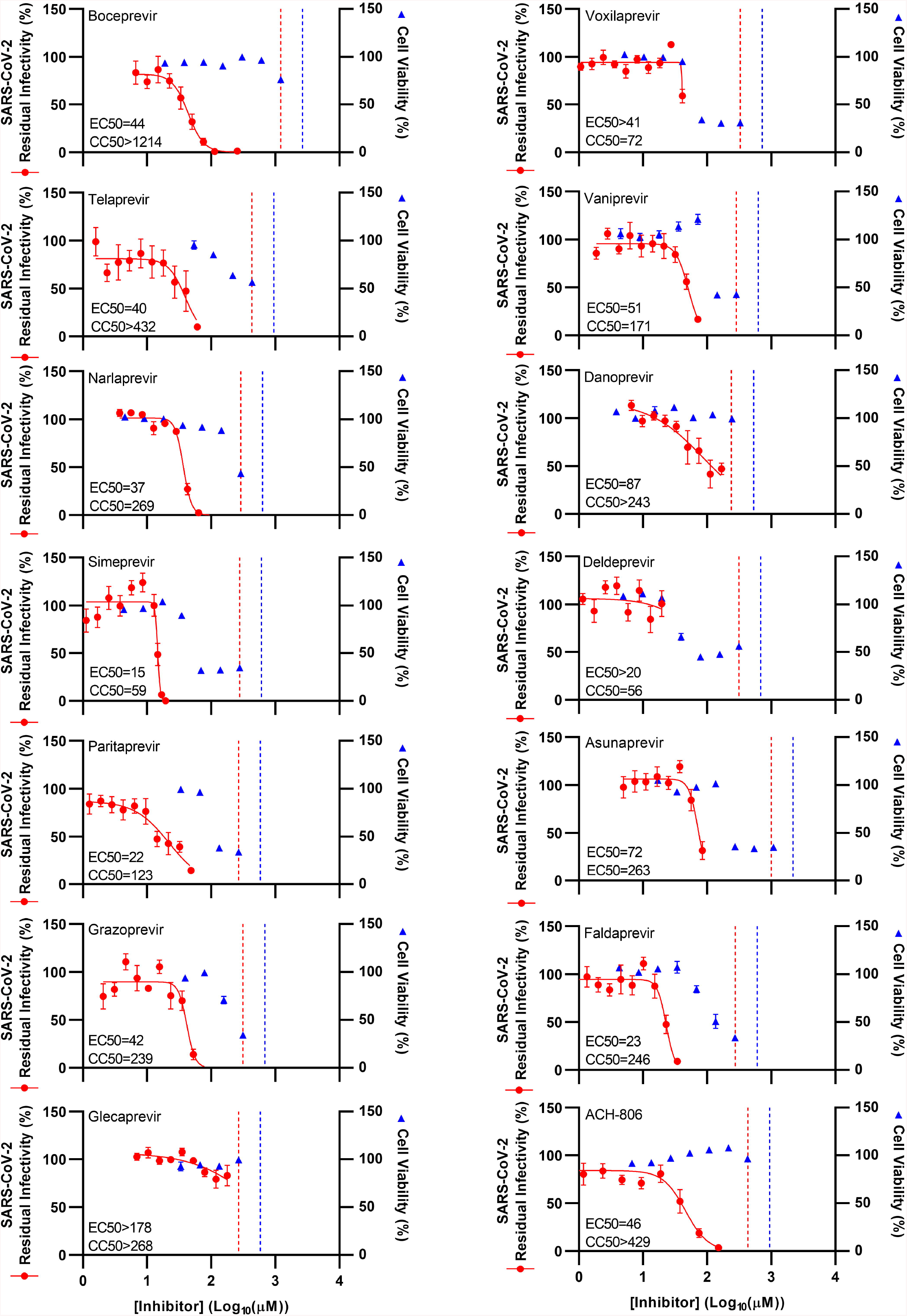
Potency of a panel of HCV PI and an HCV NS4A inhibitor against SARS-CoV-2 in VeroE6 cells. VeroE6 cells were seeded in 96-well plates and the following day infected with SARS-CoV-2 followed by treatment with specified concentrations of the PI boceprevir, telaprevir, narlaprevir, simeprevir, paritaprevir, grazoprevir, glecaprevir, voxilaprevir, vaniprevir, danoprevir, deldeprevir, asunaprevir and faldaprevir, as well as HCV NS4A inhibitor ACH-806, as described in Materials and Methods. After 46-50 hours of incubation, SARS-CoV-2 infected cells were visualized by immunostaining for the SARS-CoV-2 Spike protein and quantified by automated counting, as described in Materials and Methods. Datapoints (red dots) are means of counts from 7 replicate cultures ± standard errors of the means (SEM) and represent % residual infectivity, determined as % SARS-CoV-2 positive cells relative to means of counts from 14 replicate infected nontreated control cultures. Sigmoidal concentration response curves (red lines) were fitted and EC50 values were determined, as described in Materials and Methods. Cell viability data were obtained in replicate assays with noninfected cells using a colorimetric assay, as described in Materials and Methods. Datapoints (blue triangles) are means of 3 replicate cultures ± SEM and represent % cell viability relative to mean absorbance from 12 replicate nontreated control cultures. Sigmoidal concentration response curves were fitted and CC50 values were determined as shown in Supplementary Figure 3. The red / blue stippled line represents the drug concentrations at which DMSO is expected to induce antiviral effects with reduction of residual infectivity to <70% / cytotoxicity with reduction of cell viability to <90%, according to Supplementary Figure 2.

To confirm potency of the tested PI in human cells, selected PI were studied in similar assays in human hepatoma Huh7.5 cells. In these assays, boceprevir, simeprevir and grazoprevir showed similar concentration response curves and EC50 values as in VeroE6 cells (Figure 2, Table 1). All inhibitors were diluted in DMSO. At the applied DMSO dilutions, no antiviral effect was observed in VeroE6 and Huh7.5 cells (Supplementary Figure 2); in Figure 1 and 2, drug concentrations at which DMSO was expected to induce antiviral effects are indicated. Cell viability assays were carried out for all studied drugs to determine their level of *in vitro* cytotoxicity and CC50 values. In these assays drug concentrations were used at which no DMSO induced cytotoxicity was observed (Supplementary Figure 2); in Figure 1 and 2, drug concentrations at which DMSO was expected to induce cytotoxicity are indicated. In VeroE6 cells, the linear PI showed the lowest cytotoxicity with all CC50 values above 200 μM (>1214, >432 and 269 μM for boceprevir, telaprevir and narlaprevir, respectively) (Figure 1, Table 1, Supplementary Figure 3). Among the macrocyclic inhibitors, grazoprevir, glecaprevir and danoprevir showed the lowest cytoxicity with CC50 above 200 μM. Paritaprevir and vaniprevir showed intermediate cytotoxicity with CC50 between 100 and 200 μM, while simeprevir, voxilaprevir and deldeprevir showed the highest cytotoxicity with CC50 between 50 and 100 μM. Cell viability assays carried out in Huh7.5 cells using boceprevir, telaprevir and grazoprevir showed similar results (Figure 2, Table 1, Supplementary Figure 4).

**Figure 2.**
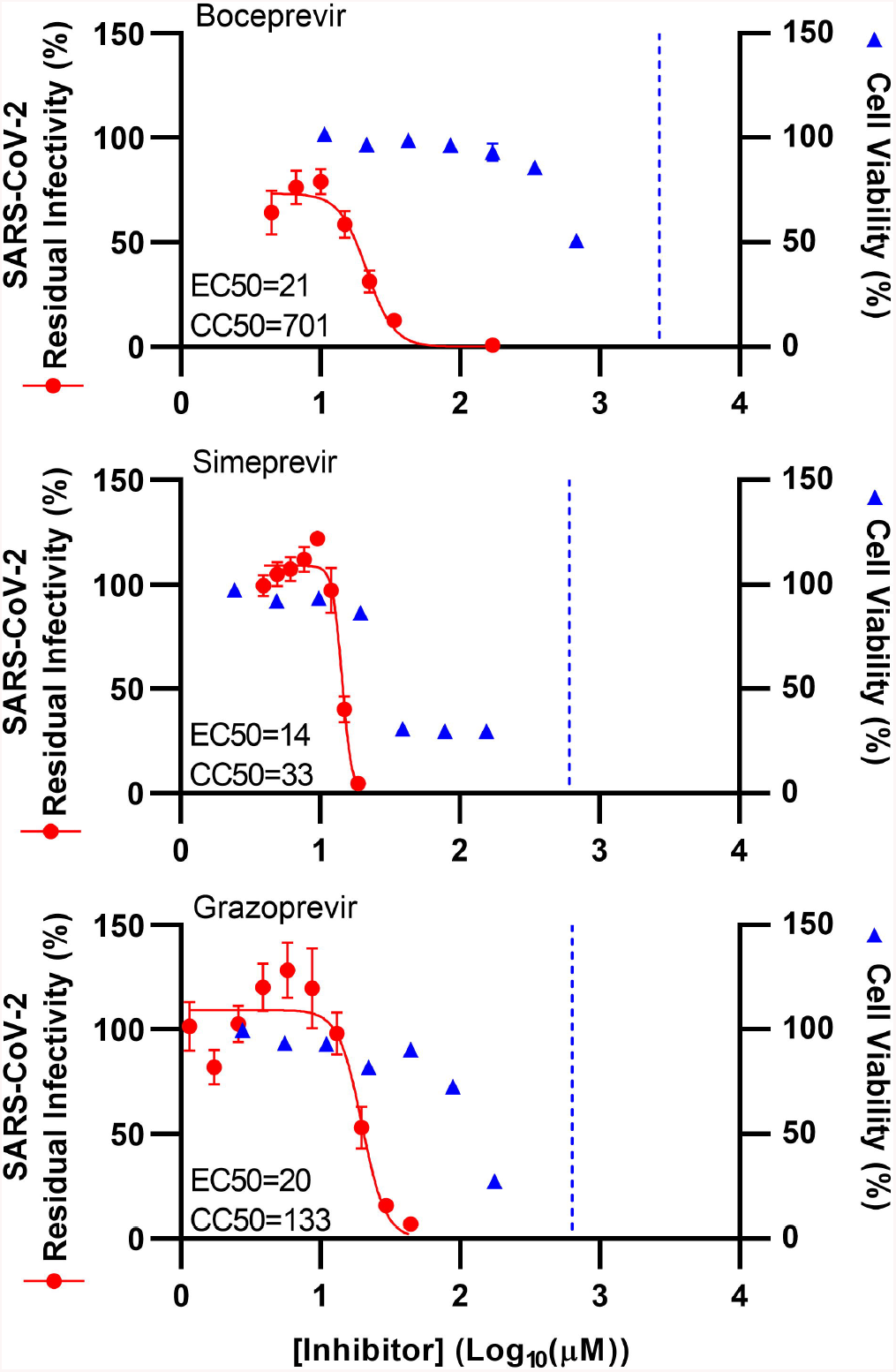
Potency of selected HCV PI against SARS-CoV-2 was confirmed in Huh7.5 cells. Huh7.5 cells were seeded in 96-well plates and the following day infected with SARS-CoV-2 followed by treatment with specified concentrations of the PI boceprevir, telaprevir and simeprevir, as described in Materials and Methods. After 70-74 hours incubation SARS-CoV-2 infected cells were visualized by immunostaining for the SARS-CoV-2 Spike protein and quantified by automated counting, as described in Materials and Methods. Datapoints (red dots) are means of 7 replicates ± SEM and represent % residual infectivity, determined as % SARS-CoV-2 positive cells relative to means of counts from 14 replicate infected nontreated control cultures. Sigmoidal concentration response curves (red lines) were fitted and EC50 values were determined, as described in Materials and Methods. Cell viability data were obtained in replicate assays with noninfected cells using a colorimetric assay as described in Materials and Methods. Data points (blue triangles) are means of 3 replicate cultures ± SEM and represent % cell viability relative to mean absorbance of 12 nontreated controls. Sigmoidal concentration response curves were fitted and CC50 values were determined, as shown in Supplementary Figure 4. The blue stippled line represents the drug concentrations at which DMSO is expected to induce cytotoxicity with reduction of cell viability to <90%, according to Supplementary Figure 2; DMSO did not induce antiviral effects in the tested dose ranges (Supplementary Figure 2).

Based on these assays, the linear inhibitors had the highest selectivity indexes (SI=CC50/EC50), >27.6 for boceprevir, >10.8 for telaprevir and 7.3 for narlaprevir (Table 1). Of the macrocyclic inhibitors, paritaprevir and grazoprevir had the highest SI (5.6 and 5.7, respectively), while simeprevir and vaniprevir had slightly lower SI of 3.9 and 3.4, respectively. For glecaprevir, voxilaprevir, danoprevir, and deldeprevir SI could not be determined. For the acyclic inhibitors, SI values were 6.3 for faldaprevir and 3.7 for asunaprevir. Finally, SI values calculated based on assays in Huh7.5 cells were comparable to those based on assays in VeroE6 cells (Table 1). For the HCV NS4A inhibitor ACH-806, in VeroE6 cells EC50 was 46 μM, CC50 was >429 μM and SI was >9.3 (Figure 1, Table 1, Supplementary Figure 3).

### HCV PI showed differential interactions with remdesivir

To study interactions between selected PI and remdesivir, 96-well based synergy assays were carried out using the method of Chou and Talalay^32,33^, as described in Materials and Methods, and as previously applied for studies on HCV.^26^ SARS-CoV-2 infected VeroE6 cells were treated with selected PI singly, in combination with remdesivir and with remdesivir alone using dilution series, which were chosen based on determined EC50 values (Table 1, Supplementary Figure 5, and using fixed concentration ratios.

For each treatment condition, the inhibition percentage was determined as a fractional effect (Fa) value. Using the CompuSyn software, Fa values were plotted against the concentrations of the single inhibitors and combinations of inhibitors, resulting in concentration-Fa curves (Figure 3A). Additionally, combination indexes (CI) were calculated for generation of Fa-CI plots and Fa-Log_10_CI plots (Figure 3B and C). CI values <0.9 suggested synergism and CI values ≥1.1 suggested antagonism, while CI values ≥0.9 and <1.1 suggested nearly additive effects; categories for the interaction level are further specified in Materials and Methods and in Figure 4. Lastly, drug reduction indexes (DRI) were calculated for generation of Fa-DRI plots and Fa-Log_10_DRI plots (Figure 3D and E). DRI values indicated how many folds the concentration of each inhibitor of the tested combination could be reduced due to synergism. Obtained CI and DRI values are summarized in Figure 4.

**Figure 3.**
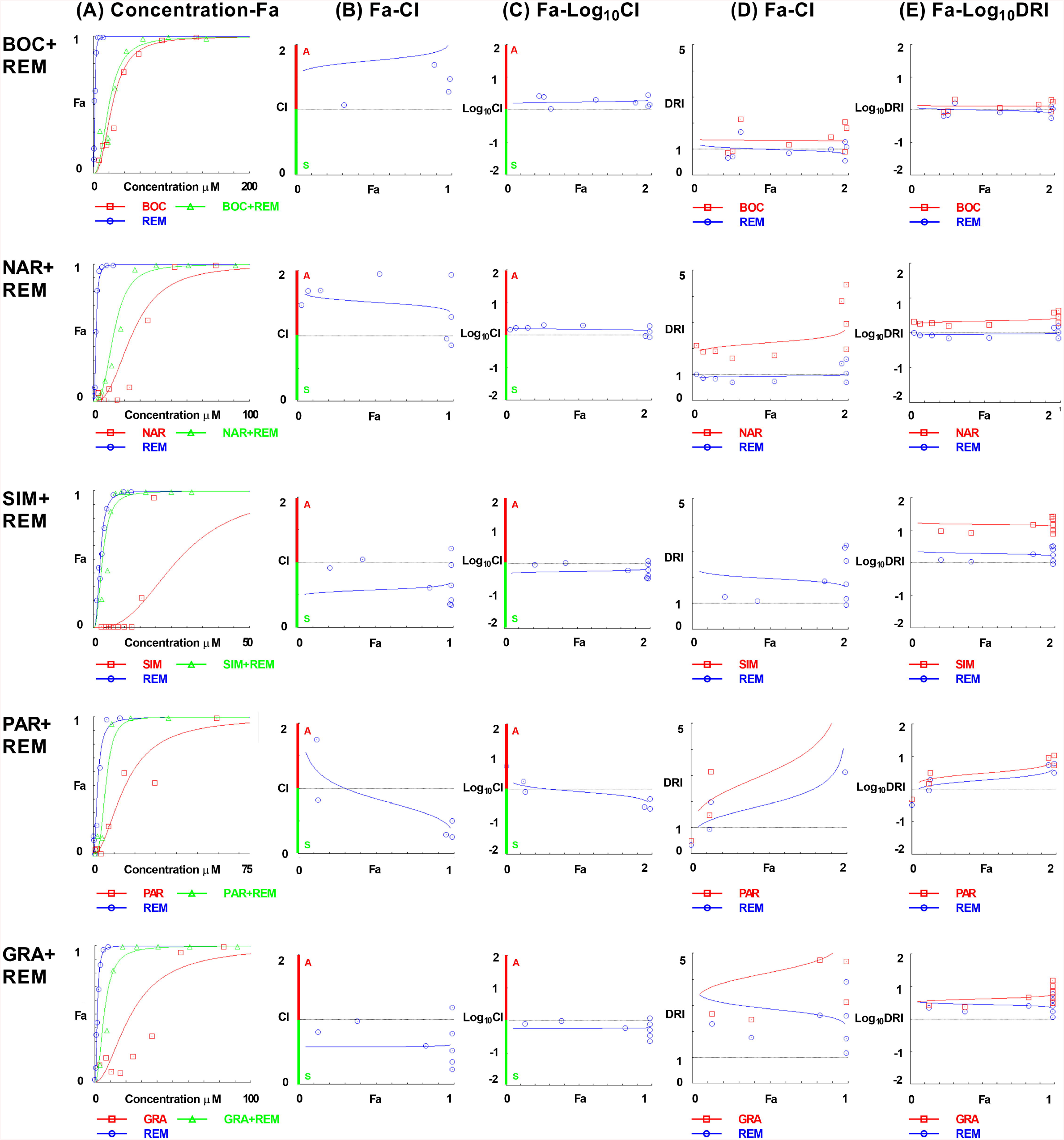
Analysis of interactions of selected HCV PI with remdesivir. VeroE6 cells seeded in 96-well plates were infected the following day with SARS-CoV-2 followed by treatment with specified concentrations of the linear PI boceprevir (BOC) and narlaprevir (NAR), or the macrocyclic PI simeprevir (SIM), paritaprevir (PAR) and grazoprevir (GRA), or polymerase inhibitor remdesivir (REM), or a combination of these PI and remdesivir, as described in Materials and Methods. After 46-50 hours incubation SARS-CoV-2 infected cells were visualized by immunostaining for SARS-CoV-2 Spike protein and quantified by automated counting, as described in Materials and Methods. Fractional effect (Fa) values were calculated by relating counts from infected and treated cultures to the mean count from at least 21 infected nontreated cultures and were entered into CompuSyn software. Datapoints are means of 6 to 7 replicates, and for each treatment experiment 6 to 10 datapoints were entered. For each inhibitor combination depicted per row, the following curves were fitted using Compusyn: (A) concentration-Fa curves plotting Fa values ranging from 0.01 to 0.99 against specified inhibitor concentrations. (B) Fa-CI curves plotting CI values ranging from 0 to 2 against Fa values ranging from 0.01 to 0.99. (C) Fa-Log_10_CI curves plotting logarithmic CI values ranging from 0.01 to 100 against Fa values ranging from 0.01 to 0.99. (B and C) Overall, CI values ≥1.1 suggest antagonism “A”, while CI values <0.9 suggest synergism “S” indicated by green and red colour coding of the y-axis, respectively. (D) Fa-DRI curves plotting DRI values ranging from 0 to 5 against Fa values ranging from 0.01 to 0.99; the datapoints and curve for simeprevir are outside the graph area and can be seen in (E). (E) Fa-Log_10_DRI curves plotting logarithmic DRI values ranging from 0.01 to 100 against Fa values ranging from 0.01 to 0.99. For an overview of CI and DRI values see Figure 4.

**Figure 4.**
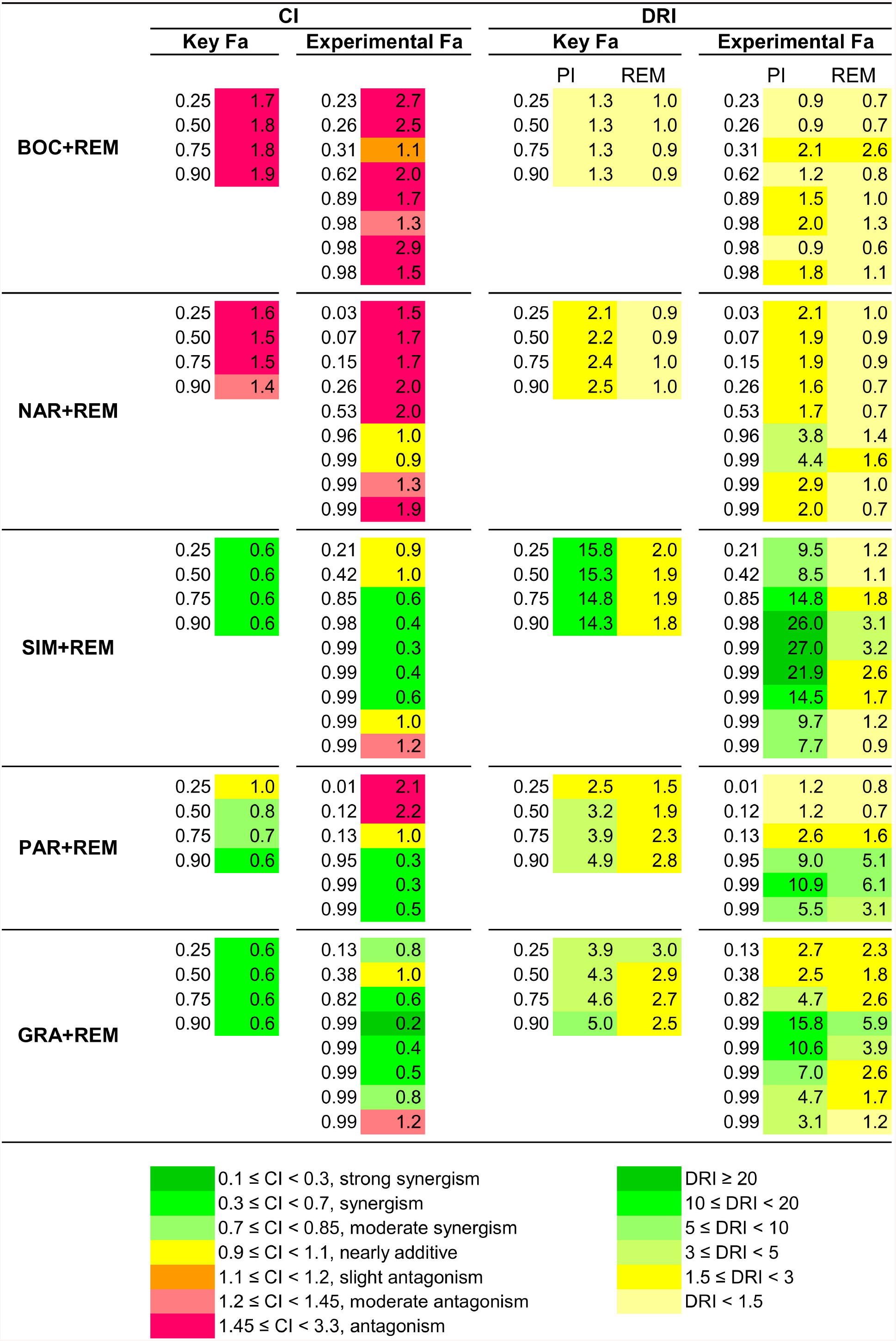
Macrocyclic versus linear PI interact with remdesivir in a synergistic versus antagonistic manner. Data are based on experiments shown in Figure 3. For the specified inhibitor combinations, boceprevir + remdesivir (BOC + REM), narlaprevir + remdesivir (NAR + REM), simeprevir + remdesivir (SIM + REM), paritaprevir + remdesivir (PAR + REM), and grazoprevir + remdesivir (GRA + REM), CI and DRI values are given. CI and DRI values at key Fa values were calculated in CompuSyn based on the curves fitted to the experimental datapoints. CI and DRI values at experimental Fa were calculated in CompuSyn. CI and DRI values are colour coded as specified, and for CI values, according to categories described in Materials and Methods; CI intervals in which no values were obtained are not included in the legend.

For the linear inhibitors boceprevir and narlaprevir, Fa-CI plots indicated at least moderate antagonism at all key Fa values, with all CI values being at least 1.4 (Figure 3B and C and Figure 4). In more detail, for boceprevir, at key Fa values antagonism was suggested by CI values of 1.7 to 1.9, and at Fa values of 0.23 to 0.98 at experimental datapoints mostly antagonism was suggested by CI values of 1.5 to 2.9. For narlaprevir, at key Fa values mostly antagonism was suggested by CI values of 1.5 and 1.6, and at Fa values 0.03 to 0.99 at experimental datapoints mostly antagonism was suggested by CI values of 1.5 to 2.0.

In line with the observed antagonism, combinations of linear PI boceprevir or narlaprevir with remdesivir showed no or only little drug reduction potential (Figure 3D and E and Figure 4). In more detail, in Fa-DRI plots DRI values for boceprevir + remdesivir were close to 1 suggesting no dose reduction potential for boceprevir or remdesivir when used in combination. For narlaprevir + remdesivir, at key Fa values DRI values for narlaprevir were 2.1 to 2.5, and at Fa values 0.03 to 0.99 from experimental datapoints DRI values were 1.6 to 4.4 suggesting a low drug reduction potential of up to 4-fold at relatively high inhibitor concentrations. DRI values for remdesivir were close to 1 suggesting no dose reduction potential.

In contrast, for the macrocyclic inhibitors simeprevir, paritaprevir and grazoprevir Fa-CI plots suggested synergism with remdesivir (Figure 3B and C and Figure 4). In more detail, for simeprevir, at key Fa values synergism was suggested by CI values of 0.6, and at Fa values of 0.21 to 0.99 from experimental datapoints, mostly synergism was suggested by CI values of 0.3 to 0.6. Similarly, for paritaprevir, at key Fa values mostly moderate synergism was suggested by CI values of 0.7 and 0.8, and at Fa values of 0.01 to 0.99 from experimental data points, mostly synergism was suggested by CI values of 0.3 and 0.5. For grazoprevir at key Fa values synergism was suggested by CI values of 0.6, and at Fa values of 0.13 to 0.99 obtained at experimental datapoints mostly synergism was suggested by CI values of 0.4 to 0.6.

In line with the observed synergism, combinations of macrocyclic PI simeprevir, paritaprevir or grazoprevir with remdesivir showed significant drug reduction potential (Figure 3D and E and Figure 4). In more detail, for simeprevir + remdesivir, at key Fa values DRI values for simeprevir were 14.3 to 15.8, and at Fa values of 0.21 to 0.99 from experimental datapoints DRI values were 7.7 to 27, suggesting a drug reduction potential of up to 27-fold for simeprevir at relatively high inhibitor concentrations. DRI for remdesivir were significantly lower, with DRI values of 1.8 to 2.0 at key Fa values and DRI values of 0.9 to 3.2 at Fa values of 0.21 to 0.99 from experimental datapoints, suggesting a drug reduction potential of up to 3-fold for remdesivir at relatively high inhibitor concentrations.

For paritaprevir + remdesivir, at key Fa values DRI values for paritaprevir were 2.5 to 4.9, and at Fa values of 0.01 to 0.99 from experimental datapoints DRI values were 1.2 to 10.9, suggesting a drug reduction potential of up to 11-fold for paritaprevir at relatively high inhibitor concentrations. DRI for remdesivir were lower, with DRI values of 1.5 to 2.8 at key Fa values and DRI values of 0.7 to 6.1 at Fa values of 0.01 to 0.99 from experimental datapoints, suggesting drug reduction potential of up to 6-fold for remdesivir at relatively high inhibitor concentrations.

Lastly, for grazoprevir + remdesivir, at key Fa values DRI values for grazoprevir were 3.9 to 5.0, and at Fa values of 0.13 to 0.99 from experimental datapoints DRI values were 2.5 to 15.8, suggesting a drug reduction potential of up to 16-fold for grazoprevir at relatively high inhibitor concentrations. DRI for remdesivir were lower, with DRI values of 2.5 to 3.0 at key Fa values and DRI values of 1.2 to 5.9 at Fa values of 0.13 to 0.99 from experimental datapoints, suggesting a drug reduction potential of up to 6-fold for remdesivir at relatively high inhibitor concentrations. DMSO used at dilutions applied in cultures treated with inhibitor combinations did not show antiviral effects in VeroE6 cells (Supplementary Figure 2). In addition, cell viability assays revealed that the applied PI + remdesivir combinations did not result in cytotoxicity in VeroE6 cells (Supplementary Figure 6).

### HCV PI showed small differences in barrier to escape of SARS-CoV-2

In order to investigate their barriers to escape, all PI except glecaprevir, voxilaprevir and deldeprevir, for which no EC50 could be determined, were used for longer-term treatment of SARS-CoV-2 infected VeroE6 cells in culture flasks at the highest possible equipotent concentration (1-fold EC50) according to predicted cytotoxicity (Figure 1, Table 1, Supplementary Figure 3). In the nontreated control cultures, infection spread to 50% of culture cells on day 1 and to 90% of culture cells on day 3 post infection, as estimated by immunostaining for the SARS-CoV-2 Spike protein (Figure 5). Following day 3, typically massive virus induced cell death was observed in these control cultures. For all PI treated cultures (Figure 5), initial viral suppression was observed with 10-30% infected culture cells on day 1 post infection and treatment initiation. On day 3, only narlaprevir, grazoprevir, vaniprevir, asunaprevir and faldaprevir treated cultures showed viral suppression with infection of 10-50% of culture cells, while in boceprevir, telaprevir, simeprevir, paritaprevir and danoprevir treated cultures 90% of culture cells were infected. On day 5, virus spread to 90% of culture cells in grazoprevir treated cultures, while cultures treated with vaniprevir and asunaprevir were closed due to massive cell death, assumed to be due to PI induced cytotoxicity, possibly enhanced by SARS-CoV-2 infection. On day 7, in narlaprevir and faldaprevir treated cultures 60% of culture cells were infected; these cultures were closed on day 9 due to massive cell death.

**Figure 5.**
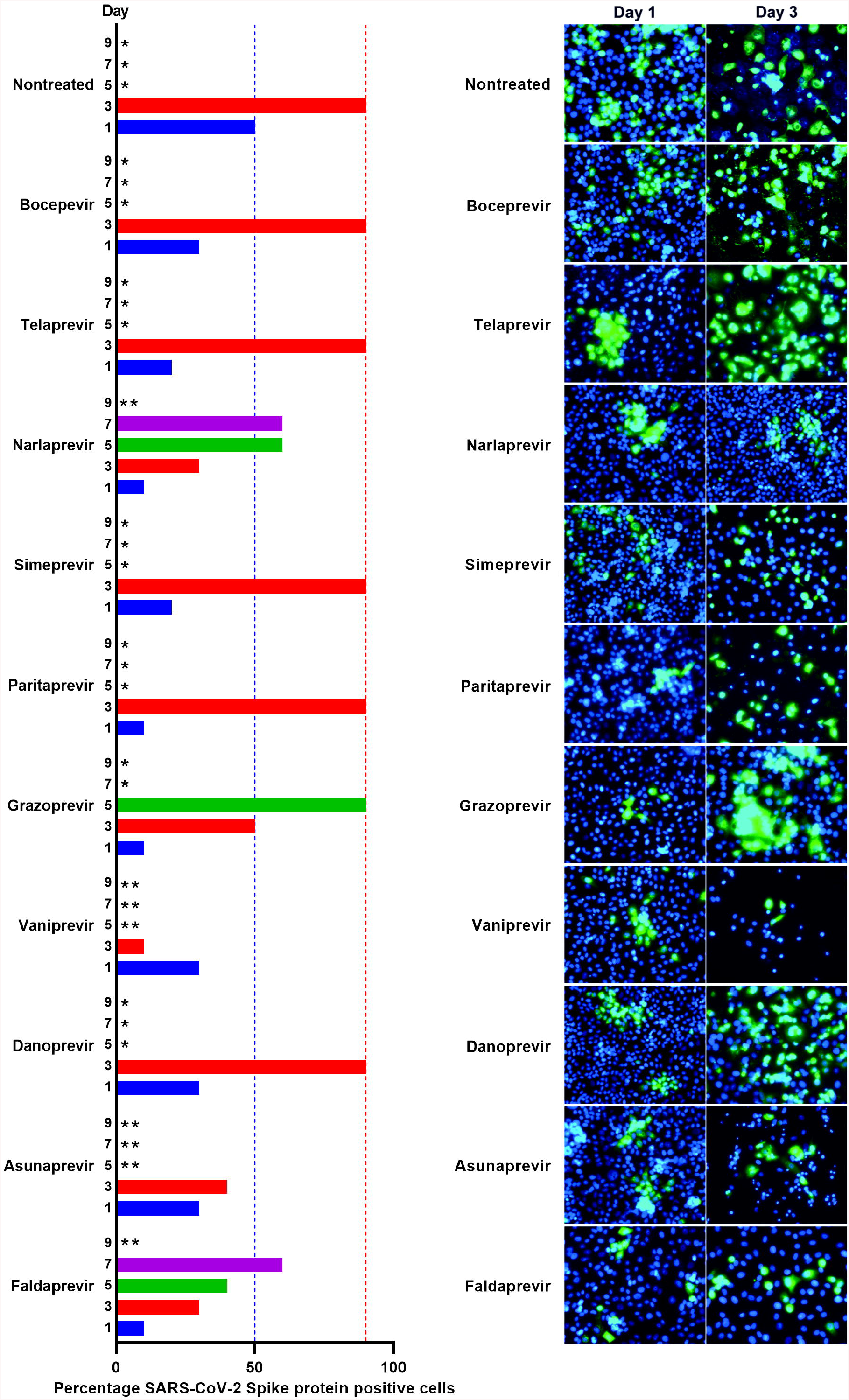
Comparison of barrier to escape for HCV PI at equipotent concentrations. VeroE6 cells seeded the previous day in T25 flasks were infected with SARS-CoV-2 followed by treatment with 1-fold EC50 of PI boceprevir, telaprevir, narlaprevir, simeprevir, paritaprevir, grazoprevir, vaniprevir, danoprevir, asunaprevir and faldaprevir, which were administered immediately after infection and on day 1, 3, 5 and 7 post infection when cells were split, as described in Materials and Methods. Left panel, the % of SARS-CoV-2 infected cells on the specified days post infection, was determined by immunostaining. Right panel, replicate cultures were derived following cell splitting and treatment and immunostained for the SARS-CoV-2 Spike protein and counterstained with Hoechst dye and images were acquired, as descried in Materials and Methods. Cultures summarized in this figure are derived from different experimental setups, each including an infected nontreated control culture, which showed viral spread comparable to that in the depicted representative culture. *Culture was terminated, or infection data not recorded, due to virus induced cell death. **Culture was terminated due to drug induced cytotoxicity, possibly enhanced by viral infection.

### Boceprevir had the potential to completely suppress viral infection *in vitro*

As suboptimal viral suppression was observed under treatment with 1-fold EC50, we chose the PI with the highest SI to enable longer-term treatment at higher fold EC50 concentrations. VeroE6 cells infected with SARS-CoV-2 were treated with 1-, 1.5-, 2-, 2.5, 3- and 5-fold EC50 of boceprevir (Figure 6). Treatment with 1- and 1.5-fold EC50 of boceprevir only had a minor impact on viral spread on day 1 post infection and treatment initiation, while 90% of culture cells became infected on day 3, as observed for nontreated control cells. Also, in cultures treated with 2 and 2.5-fold EC50, 90% of culture cells became infected on day 5. In contrast, treatment with 3- and 5-fold EC50 resulted in sustained viral suppression with no evidence of infected cells in the culture treated with 3-fold EC50 from day 3 and in the culture treated with 5-fold EC50 from day 1 during a follow up period of 7 and 17 days, respectively. In addition, from cultures treated with 3- and 5-fold EC50 on day 5 and day 3, respectively, replicate cultures receiving no treatment were derived, which did not show any infected cells during a follow up period of 10 days, suggesting that the infection was cured under these treatments.

**Figure 6.**
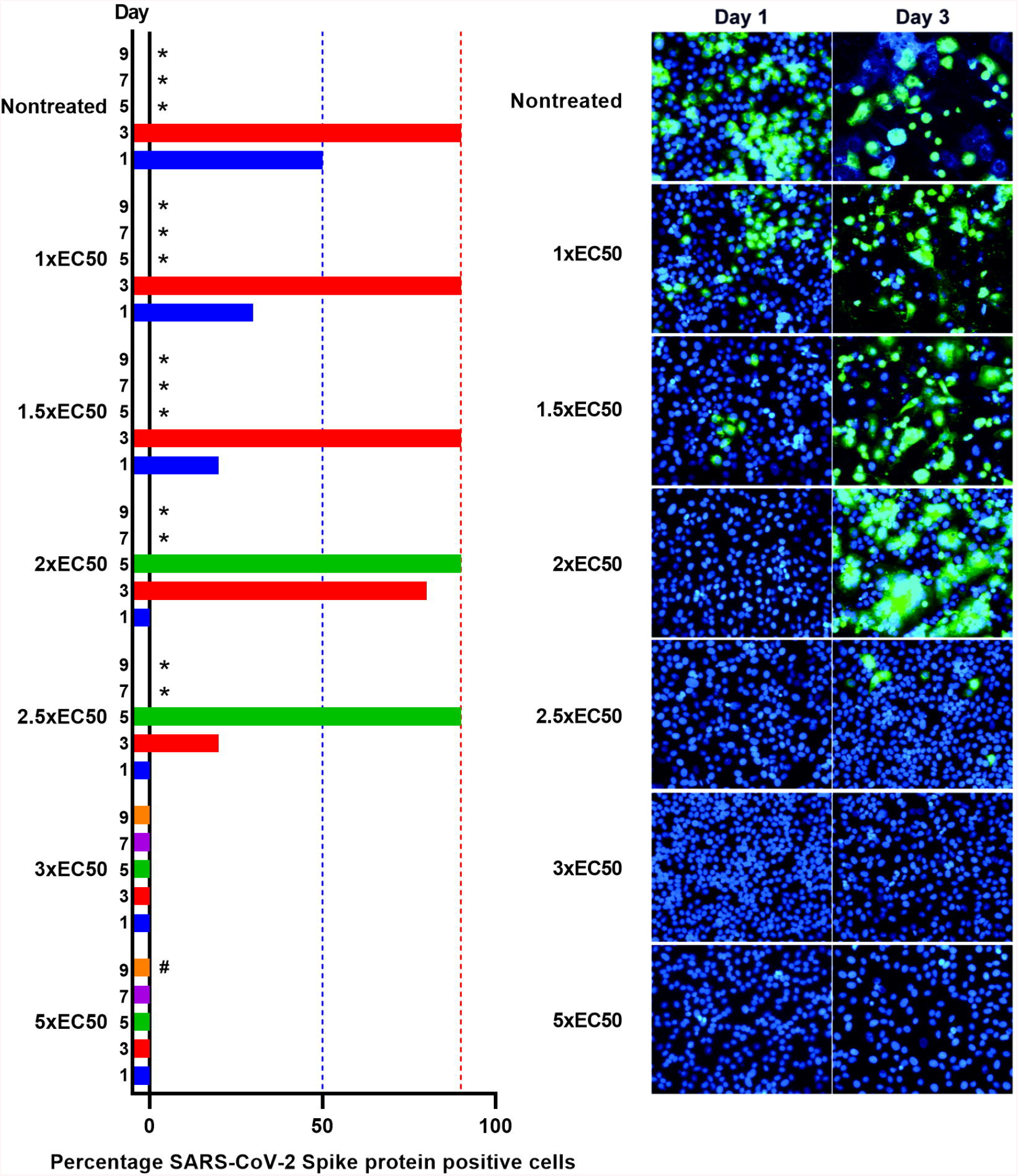
Boceprevir was capable of completely suppressing SARS-CoV-2. VeroE6 cells seeded the previous day in T25 flasks were infected with SARS-CoV-2 followed by treatment with 1-, 1.5-, 2-, 2.5-, 3- and 5-fold EC50 boceprevir, which was administered immediately after infection and subsequently at the indicated timepoints when cells were split, as described in Materials and Methods. Left panel, the % of SARS-CoV-2 infected cells on the specified days post infection was determined by immunostaining. Right panel, replicate cultures were derived following cell splitting and treatment and immunostained for the SARS-CoV-2 Spike protein and counterstained with Hoechst dye and images were acquired, as described in Materials and Methods. Cultures summarized in this figure are derived from different experimental setups, each including an infected nontreated control culture, which showed viral spread comparable to that in the depicted representative culture. *Culture was terminated, or infection data not recorded, due to virus induced cell death. ** Culture was maintained for a total of 17 days without indication of infection (no observation of single SARS-CoV-2 Spike protein positive cells).

### Simeprevir in combination with remdesivir completely suppressed viral infection *in vitro*

To further study and confirm the interactions of PI with remdesivir, 2 PI with apparent differential interactions with remdesivir were selected for longer-term treatment of SARS-CoV-2. VeroE6 cells infected with SARS-CoV-2 were treated with the PI boceprevir or simeprevir singly, remdesivir singly or either PI in combination with remdesivir. Inhibitor concentrations were selected to confer suboptimal effects in order to rule out viral suppression by treatment with single inhibitors.

Equipotent concentrations of PI were used based on data shown in Figures 1 and 5. For remdesivir, potency was evaluated based on concentration response curves obtained from data shown in Figure 3 and Supplementary Figure 5, and in addition based on pilot longer-term treatment assays (data not shown). Treatment with remdesivir, boceprevir or simeprevir singly as well as treatment with boceprevir + remdesivir had none or only a minor impact on viral spread on day 1 post infection and treatment initiation, while 80 to 90% of culture cells became infected on day 3, as observed for nontreated control cells (Figure 7). In contrast, in cultures treated with simeprevir + remdesivir complete and sustained viral suppression was achieved with no evidence of infection from day 1 during a follow up period of 15 days. In addition, to confirm complete viral suppression, from this culture a replicate culture receiving no treatment was derived on day 5, which did not show any infected cells during a follow up period of 19 days.

**Figure 7.**
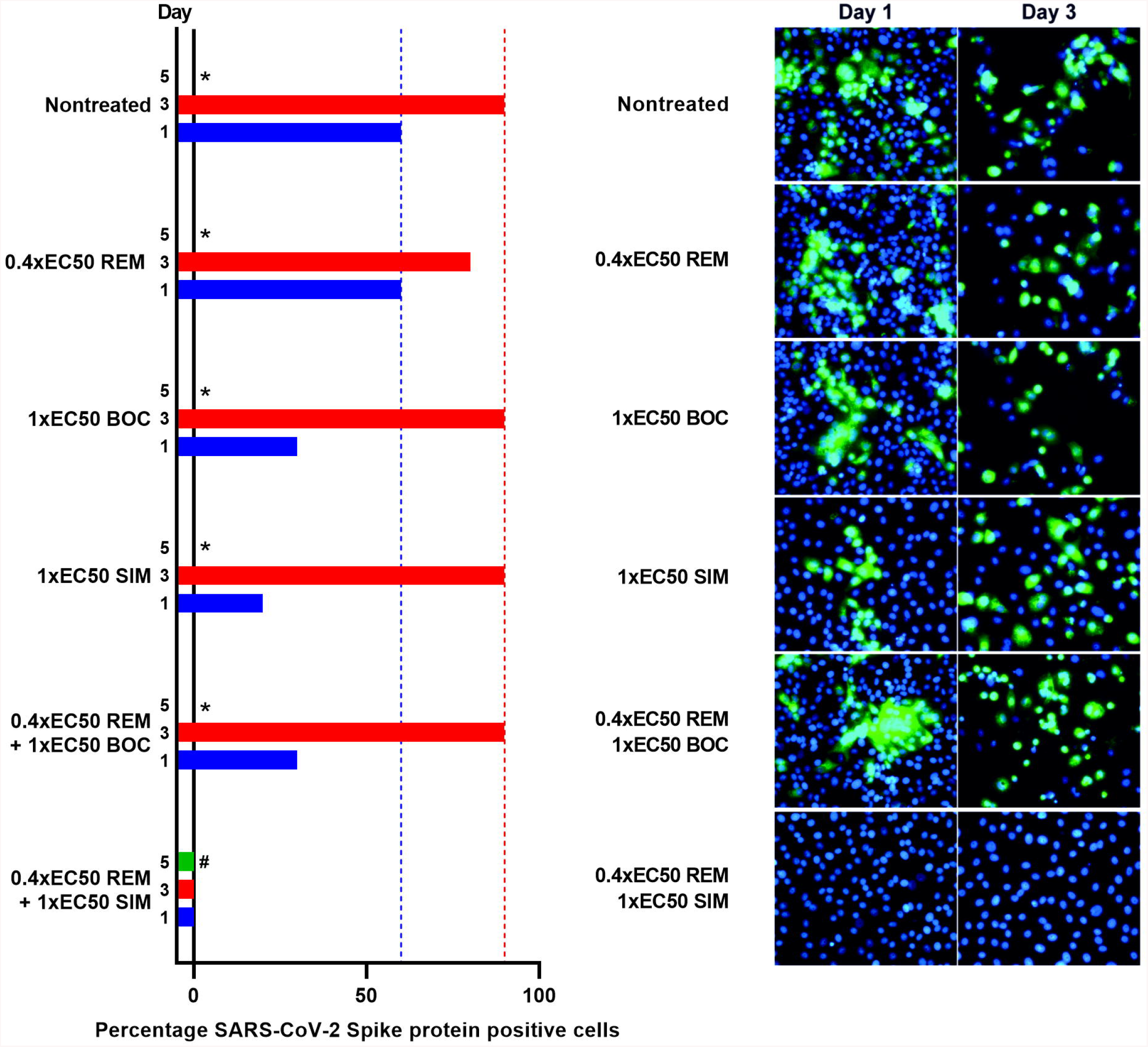
At equipotent concentrations, simeprevir but not boceprevir synergized with remdesivir to completely suppress viral infection. VeroE6 cells seeded the previous day in T25 flasks were infected with SARS-CoV-2 followed by treatment with 0.4-fold EC50 of remdesivir (REM) or 1-fold EC50 of PI boceprevir (BOC) or simeprevir (SIM), or a combination of remdesivir with either PI, including an infected, nontreated culture serving as a positive control for viral spread, as described in Materials and Methods. Treatment was administered immediately after infection and subsequently at the indicated timepoints when cells were split, as described in Materials and Methods. Left panel, the % of SARS-CoV-2 infected cells on the specified days post infection was determined by immunostaining. Right panel, replicate cultures were derived following cell splitting and treatment and immunostained for the SARS-CoV-2 Spike protein and counterstained with Hoechst dye and images were acquired, as described in Materials and Methods. *Culture was terminated, or infection data not recorded, due to virus induced cell death. ^#^Culture was maintained for a total of 15 days without indication of infection (no observation of single SARS-CoV-2 Spike protein positive cells).

## Discussion

We provided a head-to-head comparison of the efficacy of a panel of clinically relevant HCV PI, including all approved PI, against SARS-CoV-2 in cell-based assays. In short-term antiviral assays, PI showed differential potency with EC50 values between 15 μM (simeprevir) and >178 μM (glecaprevir). Detailed short-term synergy studies with a PI sub-panel revealed PI structure dependent interactions with remdesivir, with linear inhibitors showing antagonism and macrocyclic inhibitors showing synergism. In longer-term cultures, at relatively low equipotent concentrations PI showed small differences regarding barrier to escape. For boceprevir, a relatively high SI facilitated treatment with higher concentrations revealing its potential to completely suppress viral infection. Further, combination of simeprevir and remdesivir suppressed viral infection at relatively low inhibitor concentrations, shown to be subtherapeutic in single treatments.

EC50 against SARS-CoV-2 were in the micromolar range, with the lowest EC50 (15 μM for simeprevir) approaching EC50 of remdesivir, determined to be 2.5 μM in this study, which was in line with previously reported results.^34,35^ However, EC50 of PI against SARS-CoV-2 were higher than EC50 against HCV: Initially developed HCV PI such as boceprevir and simeprevir were ~10- to 1,000-fold less potent, while optimized HCV PI such as grazoprevir, glecaprevir and voxilaprevir were ̴ 1,000- to 100,000-fold less potent against SARS-CoV-2 than against different HCV isolates.^24,26–28,30,36–38^ Boceprevir showed the highest SI of >27.6, while simeprevir showed one of the lowest SI of 3.9. Of note, some clinically relevant drugs such as digoxin have low therapeutic breadth with SI as low as 2,^39^ and simeprevir has proven safe in clinical practice. However, to estimate the clinical potential of inhibitors, comparison of their EC50 with clinically achievable plasma and tissue concentrations is more relevant than comparison with *in vitro* CC50 values. For most HCV PI, peak plasma concentrations (Cmax) were significantly lower than the determined EC50 values (Supplementary Table 2). The most favorable Cmax/EC50 ratio was found for simeprevir (Cmax/EC50 of ~1), followed by faldaprevir (Cmax/EC50 of ~0.2), as well as boceprevir, telaprevir and vaniprevir (Cmax/EC50 of ~0.1). For HCV PI detailed information on liver concentrations is available and in general these are higher than plasma concentrations: For faldaprevir, boceprevir, telaprevir and vaniprevir liver concentrations are ~32-, ~30-, ~35-, and 20- to 280-fold higher than plasma concentrations, and thus ~6-, ~2.3-. 4.6-, and up to 36-fold higher than EC50 values, respectively.^31,40,41^ For simeprevir, liver concentrations are 20- to 40-fold higher than plasma concentrations and thus up to 39-fold higher than EC50 values.^42^ In rats, following a single oral administration of simeprevir, concentrations in the intestine, which is permissive to SARS-CoV-2 infection,^43^ were up to 128-fold higher than in the plasma, while concentrations in other tissues were roughly equal to plasma concentrations.^44^ To further estimate the clinical potential of simeprevir for treatment of SARS-CoV-2 it would be relevant to determine concentrations in relevant tissues such as in the lungs and the kidneys in humans following multiple doses in steady state.^45^

At clinically achievable tissue concentrations, inhibitor efficacy can be improved by combination treatments with synergistic and thus drug saving effects as reported here for the macrocyclic PI simeprevir, paritaprevir and grazoprevir in combination with remdesivir and as recently suggested by Lo et al. for simeprevir in combination with remdesivir, providing a less detailed analysis than in this study.^46^ Of note, Hu et al. recently reported additive effects of boceprevir and remdesivir in short-term assays based on determination of cytopathic effects.^47^ However, our extensive results in short-term assays based on detection of viral protein followed by application of a highly relevant method for synergy evaluation, clearly demonstrated that the mode of interaction between PI and remdesivir depended on the PI structure. Thus, the linear PI boceprevir and narlaprevir showed mostly antagonism, while macrocyclic PI mostly showed synergism with up to 27-fold drug reduction effects for simeprevir. We confirmed these PI structure dependent interactions with remdesivir in a longer-term treatment assay where combination of boceprevir with remdesivir did not result in additive or synergistic effects, while for combination of simeprevir and remdesivir synergistic effects were observed. This structure dependence remains to be explained, however, it could be caused by differential cellular uptake or metabolization. Further, simeprevir might target the SARS-CoV-2 polymerase in addition to M^pro^, potentially by a different mechanism than remdesivir, which might explain the observed synergy.^46^ Finally, PI were suggested to affect innate immunity, as well as alternative viral targets, which could provide the basis for the observed differences in interactions with remdesivir.^48–68^

While remdesivir is the only antiviral directly targeting SARS-CoV-2 that is approved for treatment of patients with COVID-19, its clinical efficacy has recently been questioned.^9,69–73^ Relatively limited clinical efficacy compared to high in vitro efficacy of remdesivir^34,35^ might be due to its poor distribution to the lungs.^45^ Improving drug formulation including development of inhalable formulations and combination with simeprevir might have the potential to increase clinical efficacy of remdesivir.

While carrying out this study, several research articles addressing a potential effect of HCV PI on SARS-CoV-2 were published. Using *in silico* modelling approaches, at least 21 studies predicted binding of different HCV PI to SARS-CoV-2 M^pro^.^48–60,62–68,74^ In several of these studies HCV PI were identified as top candidates following screening of large drug libraries, however, some of these studies also yielded contradictory results. Of these studies, 10 identified simeprevir^50,53,58,60,62,63,65–68^, 5 paritaprevir^49,53,55,63,66^, 3 glecaprevir^54–56^, telaprevir^48,53,64^, boceprevir^48,57,59^ and faldaprevir^48,60,74^, 2 danoprevir^48,51^ and asunaprevir^48,60^ and 1 sovaprevir^48^, vaniprevir^48^, narlaparevir^48^ and grazoprevir^52^ to bind M^pro^. Recently, 4 groups reported studies of HCV PI voxilaprevir, boceprevir and simeprevir in cell-based antiviral assays. In line with our findings for a different SARS-CoV-2 strain, voxilaprevir did not have significant antiviral effects,^75^ while boceprevir^47,59,76^ and simeprevir^46^ had antiviral effects: In VeroE6 cells, boceprevir inhibited the SARS-CoV-2 strain USA-WA1/2020 with EC50 of 1-2 μM^76^ and the Wuhan strain with EC50 of 16 μM^59^, while simeprevir inhibited the BetaCoV/Hong Kong/VM20001061/2020 strain with EC50 of 4 μM.^46^ While EC50 reported in these studies and our study are in the same range, slightly higher EC50 values observed in our study are most likely caused by differences in experimental assay conditions. In this study, we used a recently developed treatment assay based on quantification of the SARS-CoV-2 Spike protein expressing cells detected by immunostaining, while other groups used assays based on quantification of viral RNA copies, virus induced cytopathogenic effects, or reduction of viral yields. Importantly, we recorded similar EC50 values in monkey VeroE6 cells and human Huh7.5 cells validating VeroE6 cells for the study of HCV PI. HCV PI were designed and optimized to bind the HCV NS3 protease. In modelling studies, 3D similarity between the HCV NS3 protease and SARS-CoV-2 M^pro^ were reported^48,60^ even though sequence homology is low. While both viral proteases are chymotrypsin like proteases, the HCV NS3 protease has a larger and more shallow binding groove.^59^ Boceprevir was confirmed to target M^pro^, as it inhibited M^pro^ in enzymatic assays^59,76^ and the M^pro^ - boceprevir crystal structure was solved.^59,77^ Simeprevir also inhibited M^pro^ in enzymatic assays^46,76^, however, with relatively low efficacy.^46,59^ Based on this finding, Lo et al. investigated alternative targets and found simeprevir to also inhibit the SARS-CoV-2 polymerase in enzymatic assays.^46^ Thus, simeprevir might have a dual mechanism of action. It should be noted that simeprevir was also described to impact cellular innate immune responses^61^ and that alternative viral targets, including nsp3 (papain like protease domain), nsp12 (polymerase), nsp13 (helicase), nsp14 (exonuclease and methyltransferase), nsp15 (endoribonuclease), nsp16 (2’-o-ribose methyltransferase), as well as structural proteins N (capsid) and Spike, were suggested for paritaprevir, grazoprevir and simeprevir by modelling studies.^53,67,78–82^ Future detailed molecular studies are required to fully define the viral targets of different HCV PI.

Finally, we report antiviral activity of the HCV NS4A inhibitor ACH-806. Future studies are required to define the SARS-CoV-2 target of this compound and to investigate its potential to inform design of SARS-CoV-2 inhibitors.

In conclusion, following initial partially contradictory reports suggesting efficacy of selected HCV PI against SARS-CoV-2, we here provide a head-to-head comparison of the efficacy of a panel of clinically relevant HCV PI against SARS-CoV-2, including detailed studies of interaction with remdesivir. Given its favorable potency, its comparatively high human plasma and tissue concentrations, and its observed synergy with remdesivir, simeprevir might be a lead candidate for design of optimized coronavirus inhibitors and might have direct clinical relevance.

## Materials and Methods

### Cell cultivation

African green monkey kidney VeroE6 cells (gift from J. Dubuisson) and human hepatoma Huh7.5 cells^83^ were maintained at 37°C and 5% CO_2_ in Dulbecco’s Modified Eagle Medium (Invitrogen, Paisley, UK) supplemented with 10% heat inactivated fetal bovine serum (Sigma, Saint Louis Missouri, USA) and 100 U/mL penicillin with 100μL streptomycin (Gibco/Invitrogen corporation, Carlsbad, California, USA). Cells were split every 2-3 days with trypsin (Sigma, Saint Louis, Missouri, USA) to maintain a subconfluent monolayer.

### Virus isolate

The corona virus isolate SARS-CoV-2/human/Denmark/DK-AHH1/2020 was derived from a swab sample from a Danish patient that was passaged in VeroE6 cells. For the experiments presented here we used a sequence confirmed 2^nd^ viral passage stock with an infectivity titer of 5.5 log_10_ TCID50/mL.^34^

### Inhibitors

All inhibitors were purchased from Acme Bioscience (Palo Alto, California, USA) and dissolved in DMSO (Sigma, Saint Louis, Missouri, USA).

### Concentration-response antiviral treatment assays for evaluation of inhibitor potency

96-well-based short-term antiviral treatment assays in VeroE6 cells and Huh7.5 cells were developed based on assays previously established for determination of the potency of HCV PI against HCV.^24–31^

### Concentration-response antiviral treatment assays in VeroE6 cells

VeroE6 cells were seeded at 10,000 cells per well in 96-well flat-bottom plates (Thermo Fischer Scientific, Roskilde, Denmark). The following day medium was changed to 50 μL fresh medium and cells were inoculated with SARS-CoV-2/human/Denmark/DK-AHH1/2020 at MOI 0.0016 by adding 50 μL virus stock diluted in medium to each well. Following 1-hour incubation at 37°C and 5% CO_2_, infected cells were treated with a dilution series of inhibitors by adding 50 μL medium with inhibitor resulting in the specified concentrations. Alternatively, cells were treated with a dilution series of DMSO alone serving as a control for antiviral activity of DMSO. All concentrations of inhibitor were tested in 7 replicates; 14 infected and nontreated and 12 noninfected and nontreated replicates were included in each assay. After 46-50 hours incubation at 37°C and 5% CO_2_, cells were subjected to immunostaining for the SARS-CoV-2 Spike protein and evaluated as described below.

### Concentration-response antiviral treatment assays in Huh7.5 cells

Huh7.5 cells were seeded at 9,000 cells per well in 96-well flat-bottom plates (Thermo Fischer Scientific). The following day cells were inoculated with SARS-CoV-2/human/Denmark/DK-AHH1/2020 at MOI 0.0198 (based on the infectivity titer determined in VeroE6 cells) by exchanging the medium with 50 μL virus stock diluted in medium. After 1-hour incubation, infected cells were treated with a dilution series of inhibitor by adding 50 μL medium with inhibitor resulting in the specified concentrations. Alternatively, cells were treated with a dilution series of DMSO alone. After 70 to 74 hours incubation, cells were subjected to immunostaining for the SARS-CoV-2 Spike protein and evaluated as described below.

### Immunostaining and evaluation of 96-well plates

Cells were fixed and virus was inactivated by submersion into methanol (J.T.Baker, Gliwice, Poland) for 20 minutes at room temperature. For immunostaining for the SARS-CoV-2 Spike protein, plates were washed 2 times with PBS-tween [PBS (Sigma, Gillingham, UK) containing 0.1% Tween-20 (Sigma, Saint Louis, Missouri)]. Then, endogenous peroxidase activity was blocked by adding H2O2 and incubating for 10 minutes followed by 2 more washes with PBS-tween and blocking by PBSK [PBS containing 1% bovine serum albumin (Roche, Mannheim, Germany) and 0.2% skim milk (Easis, Aarhus, Denmark)] for 30 minutes. Next, plates were emptied and incubated with primary antibody SARS-CoV-2 spike chimeric monoclonal antibody (Sino Biological #40150-D004, Beijing, China) diluted 1:5,000 in PBSK for 2 hours at room temperature. Then plates were washed 2 times with PBS-tween and incubated for 1 hour at room temperature with secondary antibody F(ab’)2-Goat anti-human IgG Fc Cross-Adsorbed Secondary Antibody, HRP (Invitrogen#A24476, Carlsbad, CA, USA) or Goat F(ab’)2 Anti-Human IgG – Fc (HRP), preadsorbed (Abcamab#98595, Cambridge, UK), diluted 1:2,000 in PBSK. Finally, plates were washed 2 times with PBS-tween, and SARS-CoV-2 Spike protein positive cells were stained using DAB substrate BrightDAB kit (Immunologic # BS04-110, Duiven, Netherlands) following the manufacturer’s guidelines. Plates were evaluated by automated counting of single SARS-CoV-2 Spike protein positive cells using an ImmunoSpot series 5 UV Analyzer (CTL Europe GmbH, Bonn, Germany).^84^ The mean of counts from noninfected and nontreated wells, which was usually <50, was subtracted from counts of infected wells. Counts from infected and treated wells were related to the mean count of the 14-replicate infected nontreated wells to calculate % residual infectivity; mean counts of infected nontreated wells were 3,000-4,000 for VeroE6 cells and 1,000-2,000 for Huh7.5 cells. Datapoints are given as means of 7 replicates with SEM. Sigmoidal concentration response curves were fitted and EC50 values calculated as described previously using Graphpad Prism 8.0.0 with a bottom constraint of 0 applying the formula Y= Top/(1+10^(Log10EC50-X)*HillSlope^).^31,85^ Representative images from concentration-response antiviral treatment assays are shown by Gilmore, Zhou et al.^86^

### Analysis of interactions of PI and remdesivir

Synergy versus antagonism of selected PI in combination with remdesivir for inhibition of SARS-CoV-2 was investigated using the method of Chou and Talalay^33^ and CompuSyn freeware (ComboSyn Inc.)^32^ based on protocols previously established for HCV.^26^ The experimental setup was similar to that of the concentration-response antiviral treatment assays described above. In brief, VeroE6 cells were seeded at 10,000 cells per well in 96-well flat-bottom plates, medium was changed to 50 μL fresh medium, and cells were inoculated with SARS-CoV-2/human/Denmark/DK-AHH1/2020 at MOI 0.0016 by adding 50 μL virus stock diluted in medium to each well. Following 1-hour incubation at 37°C and 5% CO_2_, infected cells were treated with a dilution series of inhibitors by adding 50 μL medium with inhibitor resulting in the specified concentrations. Alternatively, cells were treated with a dilution series of DMSO alone serving as a control for antiviral activity of DMSO. Regarding inhibitor treatment, dilution series of selected PI alone, remdesivir alone or a combination of PI and remdesivir were used that were based on EC50 values against SARS-CoV-2. Thus, for inhibitors and combinations of inhibitors 1.25- to 2-fold dilution series with at least 6 different dilutions were applied spanning the respective EC50 values aiming at achieving inhibition values ranging between 1 and 99%. For combination treatments the same PI and remdesivir concentrations as used in single treatments were applied with a fixed ratio. All treatment conditions were tested in 6 or 7 replicates including 21 to 70 infected and nontreated replicates per experiment (with at least 7 replicates per experimental plate) and 12 noninfected and nontreated replicates per experimental plate. After 46-50 hours incubation, infected cells were visualized by immunostaining for the SARS-CoV-2 Spike protein and plates were evaluated by automated counting of single SARS-CoV-2 Spike protein positive cells, as described above. Counts from infected and treated wells were related to the mean of counts from all infected nontreated wells included in the same experiment to calculate % inhibition values which were entered into CompuSyn as fractional effect (designated Fa) values, ranging from 0.01 to 0.99. For each inhibitor or inhibitor combination 6-10 datapoints were entered based on 6-10 tested concentrations. The software was then used to determine (i) concentration-effect curves for single and combination treatments, (ii) combination index (CI) values and curves in relation to Fa values, and (iii) drug reduction index (DRI) values and curves in relation to Fa values. CI values were used to grade synergism and antagonism in accordance to suggestions by CompuSyn: CI<0.1, very strong synergism; 0.1≤CI<0.3, strong synergism; 0.3≤CI<0.7, synergism; 0.7≤CI<0.85, moderate synergism; 0.85≤CI<0.9, slight synergism; 0.9≤CI<1.1, nearly additive; 1.1≤CI<1.2, slight antagonism; 1.2≤CI<1.45, moderate antagonism; 1.45≤CI<3.3, antagonism; 3.3≤CI<10, strong antagonism; CI≥10, very strong antagonism. DRI values were used to describe how many folds the concentration of each inhibitor could be reduced due to synergism during combination treatment and at given levels of inhibition, compared to each inhibitor administered alone.

### Cell viability assays

To evaluate cytotoxic effects of the inhibitors and DMSO, cell viability was monitored using the CellTiter 96 Aqueous One Solution Cell Proliferation Assay (Promega, Madison, WI, USA). VeroE6 cells and Huh7.5 cells were seeded in 96-well flat-bottom plates at 10,000 and 9,000 cells per well, respectively, and the following day treated with dilution series of inhibitors or combinations of inhibitors, by adding 100 μL of medium containing inhibitors at the specified concentrations or DMSO alone at the specified dilutions. After 46-50 hours for VeroE6 cells and after 70-74 hours for Huh7.5 cells, cell viability was evaluated following the manufacturer’s guidelines. In brief, 20 μL CellTiter 96 Aqueous One Solution Reagent was added to each well and plates were then incubated for 1 to 3 hours at 37°C and 5% CO_2_. After incubation, for each well absorbance at 492 nm was recorded by use of a FLUOstar OPTIMA 96-well plate reader (BMG, LABTECH, Offenburg, Germany). Each inhibitor concentration was tested in 2 to 4 replicate wells and each experimental plate included 12 replicate nontreated control wells. Absorbance values of treated wells were related to the mean absorbance of the nontreated wells to estimate % cell viability. Datapoints are given as means of 2 to 4 replicates with SEM. Sigmoidal dose response curves were fitted and 50% cytotoxic concentration (CC50) values were calculated using GraphPad Prism 8.0.0 with a bottom constraint of 0 applying the formula Y= Top/(1+10^(Log10EC50-X)*HillSlope)^.

### Longer-term SARS-CoV2 infected and PI treated VeroE6 cell cultures

VeroE6 cells were seeded at 10^6^ cells per flask in T25 flasks (Nunc) and the following day infected at MOI 0.000016 with SARS-CoV-2/human/Denmark/DK-AHH1/2020. Cells were treated with specified fold EC50 of inhibitors on the day of infection, by adding inhibitors together with the virus and again on day 1 post infection. Following, cells were split and treated every 2 days with the specified concentrations of inhibitors and the percentage of infected culture cells was evaluated by immunostaining for the SARS-CoV-2 Spike protein and immunofluorescence imaging, as described below. For each experiment a nontreated infected culture was included serving as a positive control for infection. Cultures were closed when massive cell death occurred, induced by viral infection and/or inhibitor treatment.

### Immunostaining and immunofluorescence imaging for evaluation of longer-term VeroE6 cell cultures

In longer-term SARS-CoV-2 infected and PI treated cultures, following cell splitting and treatment, replicate cell cultures were seeded in 8-well chamber slides (Thermo Fisher Scientific, Rochester, NY, USA). The next day, cells were fixed, and virus was inactivated by submersion into methanol for 20 minutes. Chamber slides were washed twice with PBS-tween and then incubated with primary antibody SARS-CoV-2 spike chimeric monoclonal antibody (Sino Biological #40150-D004, Beijing, China) diluted 1:1,000 in PBSK for 2 hours at room temperature. Following 2 washes with PBS-tween, chamber slides were incubated with secondary antibody Alexa-Fluor 488 goat anti-human IgG (H+L) (Invitrogen #A-11013, Paisley, UK) diluted 1:500 and Hoechst 33342 (Invitrogen, Paisley, UK) diluted 1:1,000 in PBS-tween for 20 minutes at room temperature. The percentage of SARS-CoV-2 Spike protein positive cells was evaluated by fluorescence microscopy (ZEISS Axio Vert.A1, Jena, Germany), assigning the following designations: 0% infected cells (no cells infected), single infected cells, and 10%–90% infected cells (in steps of 10%). The images were acquired with ZEN 3.0 software.

## Abbreviations

CC50: 50% cytotoxic concentration(s);
CI: combination index(es);
COVID-19: coronavirus disease 2019;
DMSO: dimethyl sulfoxide;
DRI: drug reduction index(es);
EC50: 50% effective concentration(s);
Fa: fractional effect(s);
FDA: Food and Drug Administration;
HCV: hepatitis C virus;
M^pro^: coronavirus main protease;
NS, HCV: nonstructural protein;
nsp: SARS-CoV-2 nonstructural protein;
PBS: phosphate buffered saline;
PBSK, PBS: containing 1% bovine serum albumin and 0.2% skim milk;
PI: protease inhibitor(s);
SARS-CoV-2: severe acute respiratory syndrome coronavirus 2;
SEM: standard error of the mean(s);
SI: selectivity index(es).

## Acknowledgements

This work was supported by PhD stipends from the Candys Foundation (K.A.G., A.O.) and the China Scholarship Council (Y.Z.), and by grants from the Novo Nordisk Foundation (J.B.), Weimann Foundation (U.F.), and the Danish Agency for Science and Higher Education (J.B.). We thank Dr. Bjarne Ørskov Lindhardt (Copenhagen University Hospital, Hvidovre) and Prof. Carsten Geisler (University of Copenhagen) for support from Hvidovre Hospital and the University of Copenhagen. We thank Line Abildgaard Ryberg for discussion on structural interactions of M^pro^ and PI, as well as Pia Pedersen, Lotte Mikkelsen and Anna-Louise Sørensen (Copenhagen University Hospital, Hvidovre) for general laboratory assistance. We thank Prof. Jean Dubuisson and Dr. Sandrine Belouzard for providing VeroE6 cells.

## Author Contributions

S.R., J.B. and J.M.G. conceived this project. K.A.G., Y.Z., A.G., S.R. and J.M.G. designed the experiments. K.A.G., Y.Z., A.G. and A.O. carried out the experiments. K.A.G., Y.Z. A.G. and J.M.G. analyzed and interpreted the data. K.A.G., Y.Z., A.O., L.P., U.F., S.F. and S.R. contributed to isolation and characterization of SARS-CoV-2/human/Denmark/DK-AHH1/2020 in vitro and established experimental systems. K.A.G., Y.Z. and J.M.G. prepared an initial manuscript draft. All authors contributed to and discussed the manuscript. J.M.G. supervised the study.

## Conflict of Interest

none.

## Notes

### Competing Interest Statement

The authors have declared no competing interest.

