## Supplementary Information for "Hepatitis C Virus Protease Inhibitors Show Differential Efficacy and Interactions with Remdesivir for Treatment of SARS-CoV-2 *in Vitro*"

**Running title:** Efficacy of HCV Protease Inhibitors against SARS-CoV-2

Karen A. Gammeltoft*, Yuyong Zhou*, Andrea Galli, Anna Offersgaard, Long V. Pham, Ulrik Fahnøe, Shan Feng, Santseharay Ramirez, Jens Bukh, Judith M. Gottwein#

Copenhagen Hepatitis C Program (CO-HEP), Department of Infectious Diseases, Hvidovre Hospital and Department of Immunology and Microbiology, Faculty of Health and Medical Sciences, University of Copenhagen, Copenhagen, Denmark

* These two authors contributed equally to this work

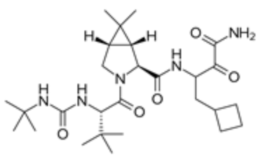

Boceprevir

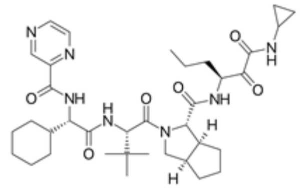

Telaprevir

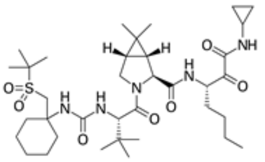

Narlaprevir

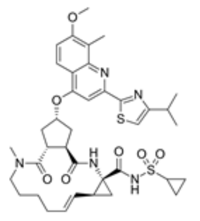

Simeprevir

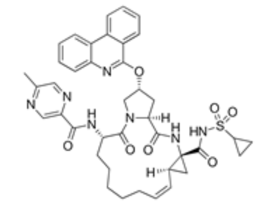

Paritaprevir

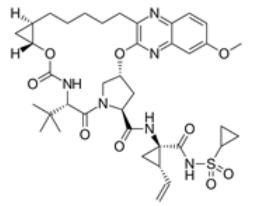

Grazoprevir

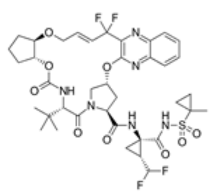

Glecaprevir

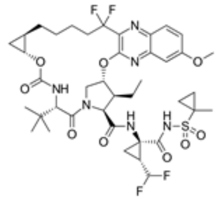

Voxilaprevir

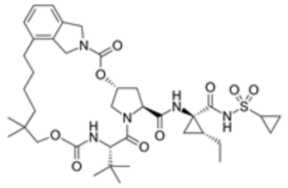

Vaniprevir

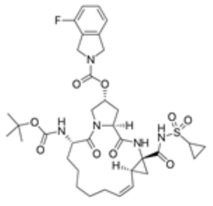

Danoprevir

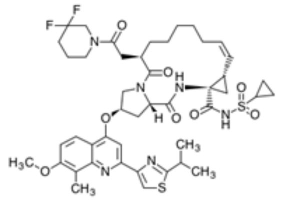

Deldeprevir

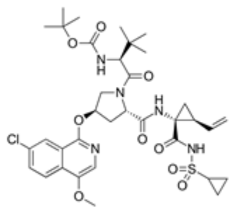

Asunaprevir

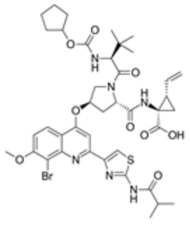

Faldaprevir

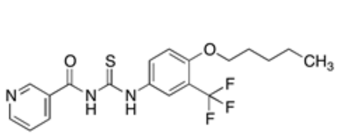

ACH-806

**Supplementary Figure 1. Structural formulas of HCV inhibitors used in this study.**

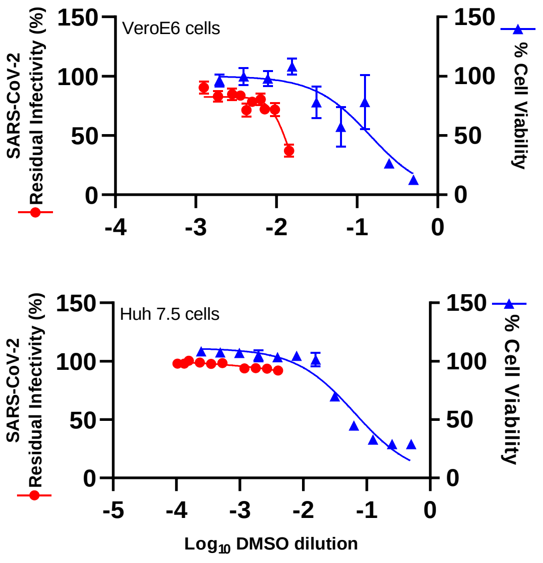


| VeroE6 cells | [Inhibitor] (µM)[a] | DMSO dilution[b] |
|---|---|---|
| **Single inhibitors** | | |
| Boceprevir | 254.9 | 333 |
| Telaprevir | 95.7 | 210 |
| Narlaprevir | 60.6 | 500 |
| Simeprevir | 19.4 | 1000 |
| Paritaprevir | 48.3 | 391 |
| Grazoprevir | 79.1 | 280 |
| Glecaprevir | 177.6 | 105 |
| Voxilaprevir | 40.9 | 560 |
| Vaniprevir | 71.4 | 280 |
| Danoprevir | 167.1 | 105 |
| Deldeprevir | 19.5 | 1120 |
| Asunaprevir | 84.2 | 833 |
| Faldapreir | 34.2 | 560 |
| ACH-806 | 150.2 | 200 |
| Remdesivir | 12.9 | 1286 |
| **Combinations of inhibitors** | | |
| Boceprevir+ Remdesivir | 182.8 | 429 |
| Narlaprevir+ Remdesivir | 91.4 | 214 |
| Simeprevir+ Remdesivir | 31.6 | 576 |
| Paritaprevir+ Remdesivir | 36.1 | 507 |
| Grazoprevir+ Remdesivir | 91.6 | 233 |
| **Huh7.5 cells** | | |
| **Single inhibitors** | | |
| Boceprevir | 170.0 | 500 |
| Simeprevir | 18.7 | 1042 |
| Grazoprevir | 44.0 | 500 |

**Supplementary Figure 2. Effect of DMSO on cell viability and antiviral effect of DMSO on SARS-CoV-2 in VeroE6 cells and Huh7.5 cells and comparison to used drug concentrations.** Left panel: Cells seeded the previous day in 96-well plates were infected with SARS-CoV-2 and treated with specified dilutions of DMSO as described in Materials and Methods. After 48 hours for VeroE6 cells and 72 hours Huh7.5 cells, cells were subjected to immunostaining for the SARS-CoV-2 Spike protein. Single infected cells were counted by automated counting and % residual infectivity was calculated by relating means of counts of infected and treated wells to the mean count of 14 replicate infected nontreated control wells. Datapoints (red dots) represent means of 7 replicates ± standard errors of the means (SEM). Sigmoidal concentration response curves were fitted using Graphpad Prism 8.0.0 and the formula Y= Top/(1+10$^{(\text{Log10EC50-X})*\text{HillSlope}}$) with a bottom constraint of 0, as described in Materials and Methods. % cell viability was determined in replicate assays with noninfected cells as described in Materials and Methods. Datapoints (blue triangles) represent means of 3 replicates ± SEM.

Right panel: Maximum concentrations of inhibitors / inhibitor combinations and corresponding DMSO dilutions used in antiviral treatment assays. [a], Maximum concentration of inhibitor used in antiviral treatment assays. For combinations of inhibitors, inhibitor concentrations reflect the combined maximum concentration of PI and remdesivir used in the specified combination treatments. [b], DMSO dilution factor at highest inhibitor concentrations used in antiviral treatment assays.

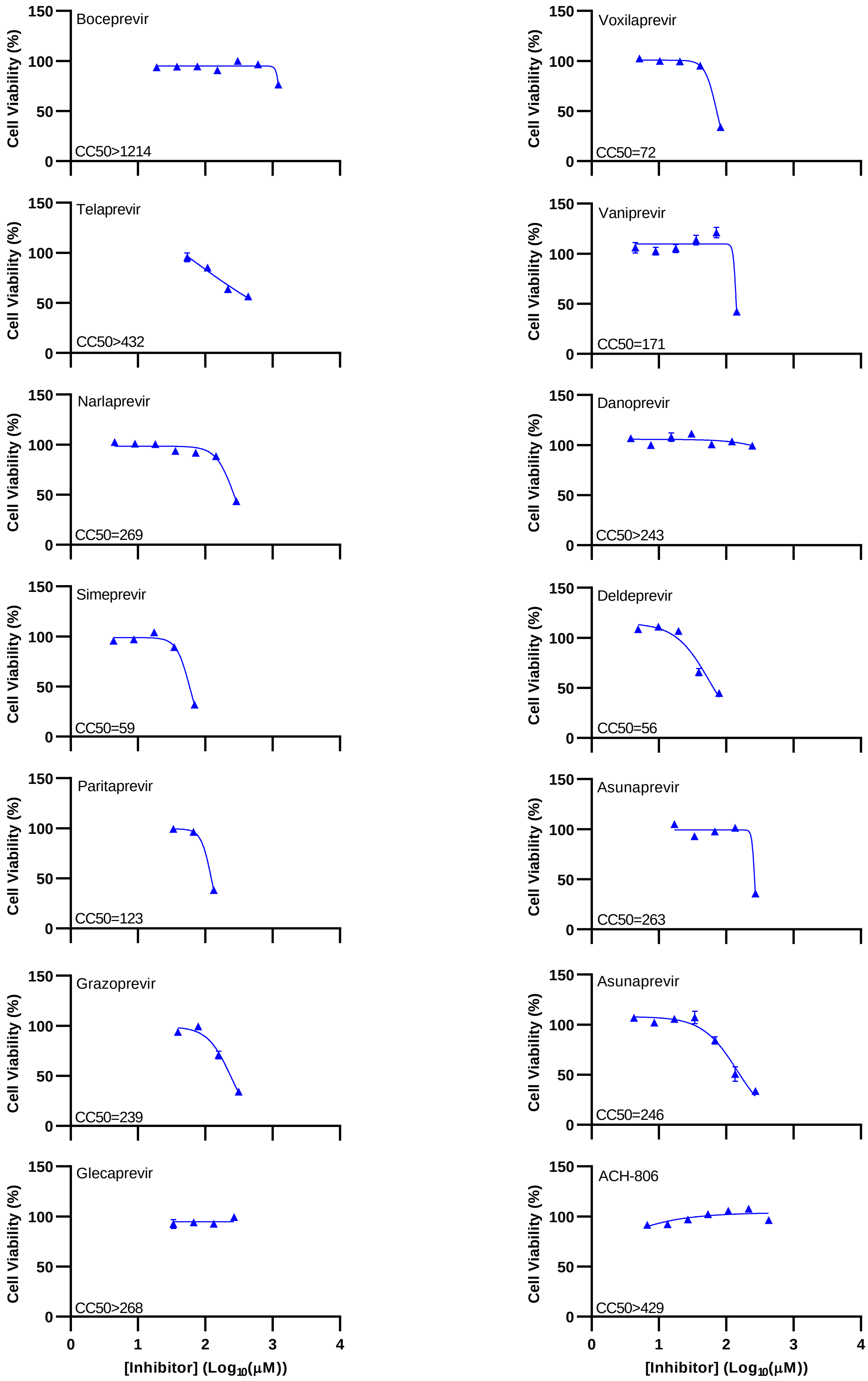
Boceprevir
CC50>1214
Voxilaprevir
CC50=72
Telaprevir
CC50>432
Vaniprevir
CC50=171
Narlaprevir
CC50=269
Danoprevir
CC50>243
Simeprevir
CC50=59
Deldeprevir
CC50=56
Paritaprevir
CC50=123
Asunaprevir
CC50=263
Grazoprevir
CC50=239
Asunaprevir
CC50=246
Glecaprevir
CC50>268
ACH-806
CC50>429
Cell Viability (%)
150
100
50
0
0
1
2
3
4
[Inhibitor] (Log10(μM))

**Supplementary Figure 3. Determination of effect of HCV PI on cell viability in VeroE6 cells and calculation of CC50 values.** 96-well-based cell viability assays were carried out for the PI boceprevir, telaprevir, narlaprevir, simeprevir, paritaprevir, grazoprevir, glecaprevir, voxilaprevir, vaniprevir, danoprevir, deldeprevir, asunaprevir, faldaprevir and for the HCV NS4A inhibitor ACH-806, as described in Materials and Methods. Sigmoidal curves were fitted and CC50 values were calculated using Graphpad Prism 8.0.0 and the formula $Y= Top/(1+10^{(Log10EC50-X)*HillSlope})$ with a bottom constraint of 0. Datapoints represent means of 3 or 4 replicates ± SEM. To facilitate curve fitting and CC50 calculation, the datapoints at the higher concentrations of the lower plateau were excluded. All datapoints generated in these experiments are shown in Figure 1.

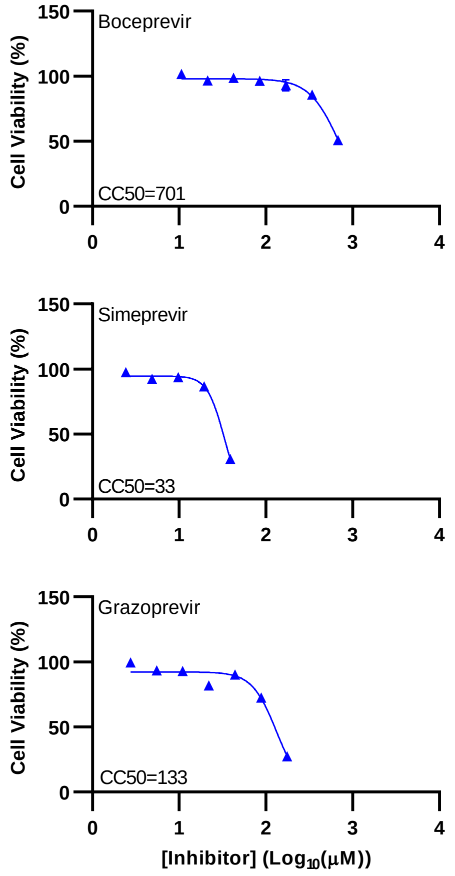


**Supplementary Figure 4. Determination of effect of selected HCV PI on cell viability in Huh7.5 cells and calculation of CC50 values.** 96-well-based cell viability assays were carried out for PI boceprevir, simeprevir and grazoprevir, as described in Materials and Methods. Sigmoidal curves were fitted and CC50 values were calculated using Graphpad Prism 8.0.0 and the formula $Y= Top/(1+10^{(Log10EC50-X)*HillSlope})$ with a bottom constraint of 0. Datapoints represent means of 3 or 4 replicates ± SEM. To facilitate curve fitting and calculation of CC50 values, the datapoints at the higher concentrations of the lower plateau were excluded. All datapoints generated in these experiments are shown in Figure 2.

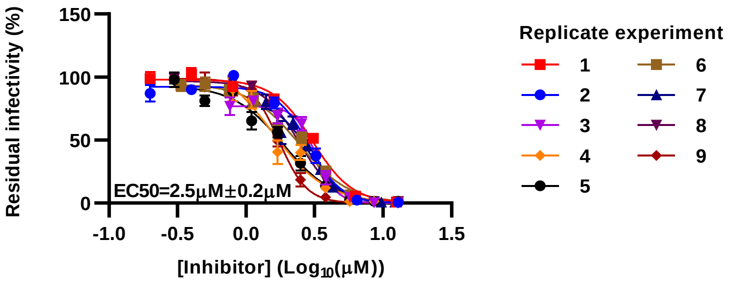


**Supplementary Figure 5. Potency of remdesivir against SARS-CoV-2 in VeroE6 cells.** VeroE6 cells seeded in 96-well plates the previous day were infected with SARS-CoV-2 followed by treatment with specified concentrations of remdesivir, as described in Materials and Methods. After 48 hours incubation SARS-CoV-2 infected cells were visualized by immunostaining for the SARS-CoV-2 Spike protein and quantified by automated counting, as described in Materials and Methods. Datapoints are means of counts from 6 to 7 replicate cultures ± SEM and represent % residual infectivity, determined as % SARS-CoV-2 positive cells relative to means of counts from 14 replicate infected nontreated control cultures. Sigmoidal concentration response curves were fitted using Graphpad Prism 8.0.0 and the formula $Y= Top/(1+10^{(Log10EC50-X)*HillSlope})$ with a bottom constraint of 0. EC50 values were determined, as described in Materials and Methods. Each curve represents a separate experiment numbered 1 to 9; the given EC50 value is the mean of EC50 values derived from the 9 experiments ± SEM. Data were generated in single treatments with remdesivir in the combination treatment experiments shown in Figure 3.

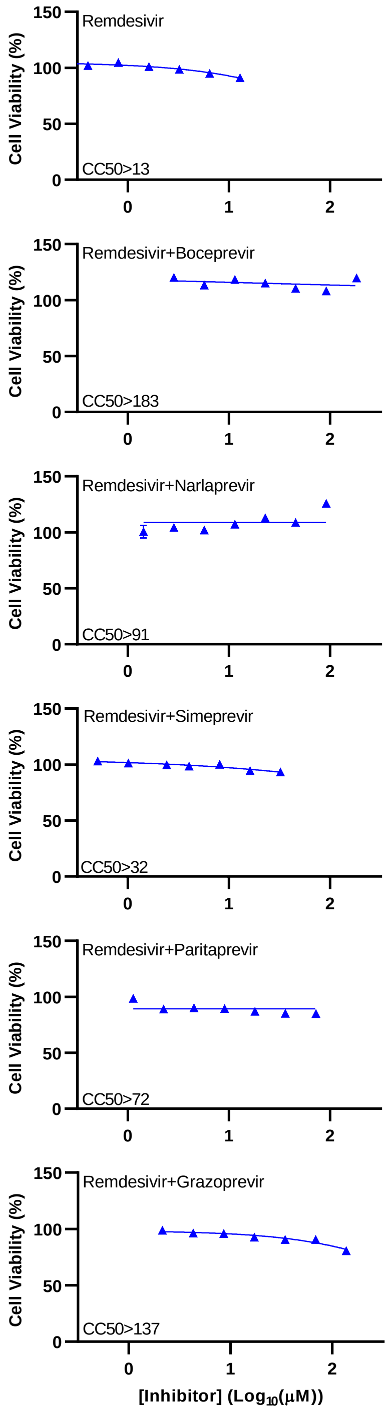


**Supplementary Figure 6. Determination of effect of remdesivir and combinations of PI and remdesivir on cell viability in VeroE6 cells.** 96-well based cell viability assays were carried out for remdesivir and PI boceprevir, narlaprevir, simeprevir, paritaprevir, grazoprevir, in combination with remdesivir, as described in Materials and Methods. Curves were fitted using Graphpad Prism

8.0.0 and the formula $Y= Top/(1+10^{(Log10EC50-X)*HillSlope})$ with a bottom constraint of 0. Datapoints represent means of 2 to 4 replicates ± SEM. Inhibitor concentrations [$Log_{10}$ µM] reflect the combined concentration of PI and remdesivir.

#### Supplementary Table 1. Overview of HCV PI and their clinical status.

| PI name | Alternative PI name | Company | Stage of clinical trials | Approvals[a] | | | | | | | | | | |
|---|---|---|---|---|---|---|---|---|---|---|---|---|---|---|
| | | | | Countries | Year | | | | | | | | | |
| | | | | | 2011 | 2012 | 2013 | 2014 | 2015 | 2016 | 2017 | 2018 | 2019 | 2020 |
| Boceprevir | none | Merck | Phase 3 | US, EU | | | | | | | | | | |
| Telaprevir | VX/950 | Janssen | Phase 3 | US, EU | | | | | | | | | | |
| Narlaprevir | SCH 900518 | Merck | Phase 3 | Russia | | | | | | | | | | |
| Simeprevir | TMC435, TMC435350 | Janssen | Phase 3 | US, EU | | | | | | | | | | |
| Paritaprevir | ABT-450 | AbbVie | Phase 3 | US, EU, China | | | | | | | | | | |
| Grazoprevir | MK-5172 | Merck | Phase 3 | US, EU, China | | | | | | | | | | |
| Glecaprevir | ABT-493 | AbbVie | Phase 3 | US, EU, China | | | | | | | | | | |
| Voxilaprevir | none | Gilead | Phase 3 | US, EU, China | | | | | | | | | | |
| Vaniprevir | MK-7009 | Merck | Phase 3 | Japan | | | | | | | | | | |
| Danoprevir | ITMN-191, RG-7227 | Roche | Phase 3 | China | | | | | | | | | | |
| Deldeprevir | ACH-2684 | Achillion | Phase 1 | Not approved | | | | | Not approved | | | | | |
| Asunaprevir | BMS-650032 | Bristol Myers Squibb | Phase 3 | Japan, Canada and China | | | | | | | | | | |
| Faldaprevir | BI 201335 | Boehringer | Phase 3 | Not approved | | | | | Not approved | | | | | |

[a]Approvals, indicating which countries / unions approved the specified PI and in which year PI were first approved (red), as well as in which year they were discontinued (peach). U.S., EU and China were considered for each PI; when PI were not approved by U.S. or EU, additional countries were considered. Several HCV PI were discontinued following initial approval as more efficient PI with increased efficacy, including efficacy against different HCV genotypes, were developed. Clinical development of deldeprevir was stopped in favor of sovaprevir also developed by Achillion, which was not available for our studies. Clinical development of faldaprevir was stopped as Boehringer exited the HCV market due to a growing number of successful HCV direct antiviral treatments becoming available.

#### Supplementary Table 2. Overview of peak plasma concentrations of HCV PI.

| PI name | $C_{max}$ (µM)[a] | Cmax/EC50[b] | Dose[c] | Combination[d] | References[e] |
|---|---|---|---|---|---|
| Boceprevir | 3.3 | 0.1 | 800mg/8hrs | | https://www.accessdata.fda.gov/drugsatfda_docs/label/2011/202258lbl.pdf |
| Telaprevir | 5.2 | 0.1 | 750mg/8hrs | | https://www.accessdata.fda.gov/drugsatfda_docs/label/2011/201917lbl.pdf |
| Narlaprevir | 1.9 | 0.05 | 100mg/24hrs | ritonavir | doi: 10.1128/AAC.01044-16c |
| Simeprevir | 14.5 | 1.0 | 200 mg/24hrs | | https://www.accessdata.fda.gov/drugsatfda_docs/nda/2013/205123Orig1s000ClinPharmR.pdf |
| Paritaprevir | 0.3 | 0.01 | 150mg/24hrs | ombitasvir, dasabuvir, ritonavir | https://www.accessdata.fda.gov/drugsatfda_docs/label/2014/206619lbl.pdf |
| Grazoprevir | 0.2 | 0.005 | 100mg/24hrs | elbasvir | https://www.accessdata.fda.gov/drugsatfda_docs/label/2017/208261s002lbl.pdf |
| Glecaprevir | 0.7 | < 0.004 | 300mg/24hrs | pibrentasvir | https://www.accessdata.fda.gov/drugsatfda_docs/label/2017/209394s000lbl.pdf |
| Voxilaprevir | 0.2 | < 0.005 | 100mg/24 hrs | velpatasvir, sofosbuvir | https://www.accessdata.fda.gov/drugsatfda_docs/label/2017/209195s000lbl.pdf |
| Vaniprevir | 6.5 | 0.1 | 850mg single dose | | doi:10.1111/cts.12482 |
| Danoprevir | 3.2 | 0.04 | 900mg/12hrs | | doi: 10.1016/S0140-6736(10)61384-0 |
| Deldeprevir | na | na | na | | na |
| Asunaprevir | 0.9 | 0.01 | 100mg/12hrs | | doi: 10.1080/14740338.2017.1397128 |
| Faldaprevir | 4.4 | 0.2 | 480mg single dose | | doi: 10.1128/AAC.03359-14 |

[a]$C_{max}$, peak plasma concentration (µM) of PI achieved in patients after administration of specified dose of PI. $C_{max}$ values were determined at steady state after multiple dosings. When this information was not available, $C_{max}$ values determined following a single dosing are reported as specified. na, not available.

[b]$C_{max}$/EC50, peak plasma concentration (µM) relative to the EC50 of inhibitor against SARS-CoV-2 in VeroE6 cells (Figure 1, Table 1).

[c]Administered dose per time that resulted in the reported $C_{max}$ values. All doses are in accordance to typically recommended clinical dosing except for simeprevir, where 150mg/24 hours is recommended, resulting in Cmax values of 5.6 µM.

[d]The reported $C_{max}$ values were determined under combination treatment, including the specified PI at the specified concentration in combination with the specified compounds.

[e]References for listed $C_{max}$ values. Values were extracted from FDA reports when available. Otherwise, $C_{max}$ values were extracted from publications on representative clinical studies.
